# Projection criteria and information risks for zero-dimensional biological dynamics across molecular, epidemic, and ecological systems

**DOI:** 10.64898/2026.08.05.743148

**Authors:** Chikoo Oosawa

## Abstract

Zero-dimensional chemical master equations, ordinary differential equations, and compartmental population models replace spatial stochastic biological systems by vectors of total counts or densities. This study asks when that projection is exact and whether information retained in spatial correlations can diagnose its practical failure. Exact Markov closure is characterized by an aggregate-rate lumpability condition: for every retained transition, the sum of microscopic transition rates must be constant over all spatial configurations with the same counts. Violations are connected to BBGKY-type correlation hierarchies and to mean-field, pair, and triplet closures. Conditional rate, finite-time predictive, memory, path-space, and correlation Kullback–Leibler risks quantify distinct losses. An exactly solvable two-compartment reaction separates structural non-closure from recovery of a well-mixed law under fast hidden mixing. Copy number and a spatial mixing–interaction ratio connect concentration, volume, diffusion, and reaction parameters to practical screening, including an Escherichia coli-scale example. The same projection logic is evaluated in controlled spatial susceptible–infectious–removed and predator–prey benchmarks. Across mixed and segregated initial conditions and four mobility regimes, pair-correlation risk was strongly associated with the error of the corresponding zero-dimensional ordinary differential equations (Spearman correlations 0.95 and 1.00; pooled 0.99). A nearest-neighbour exchange sensitivity analysis preserved the positive risk–error ranking. These benchmarks do not establish a universal threshold, but support correlation information as a transferable diagnostic for selecting among count, pair, higher-order, and explicit spatial descriptions.

## 1. Introduction

Ordinary differential equations (ODEs), chemical master equations, and compartmental population models often replace a spatially distributed biological system by a small vector of totals. Examples include biochemical copy-number models, susceptible–infectious–removed (SIR) epidemic equations, and Lotka–Volterra predator–prey equations. These reductions have been exceptionally productive, but they are not identities: molecules occupy three-dimensional cells, infections occur through local contacts, and ecological interactions occur in two– or three-dimensional habitats. Spatial stochastic models, particle reaction–diffusion methods, network epidemic models, and spatial moment methods were developed precisely because local arrangements and correlations can alter reaction, transmission, and predation rates [1–6].

The central mathematical question is whether the retained count process is autonomous. This is a problem of Markov projection or lumpability [7–9]. For stochastic reaction networks, exact species lumping and rule-based abstractions provide important reduction results [10, 11]. Those methods generally aggregate species or rules within an already well-mixed description. Here we instead ask whether spatial configurations themselves can be discarded while preserving count dynamics.

A second question arises when exact closure fails: how large is the error at the time scale and parameter regime of interest? Relative entropy and path-space information provide principled measures of stochastic coarse-graining error [12–14]. Expected Kullback–Leibler risk has also been used to decide when a local biochemical approximation is sufficiently accurate [15]. Pair and triplet approximations provide intermediate descriptions between mean-field counts and complete spatial processes in reaction kinetics, epidemics, and ecology [16–19]. Yet exact closure criteria, correlation-order diagnostics, and directly interpretable mixing parameters are rarely presented as one model-selection framework.

We use the classical aggregate-rate strong-lumpability criterion as an organizing backbone rather than claiming it as a new theorem. The contribution is its integration with spatial-to-count biological reduction, BBGKY-type correlation hierarchies, several information-risk diagnostics, measurable mixing and copy-number screens, and reproducible benchmarks at three biological scales. Pair-correlation risk tracked count-model error across molecular, epidemic, and ecological examples. We deliberately used controlled models to isolate the closure mechanism; the SIR and predator–prey examples are methodological benchmarks rather than forecasts of a particular outbreak or ecosystem.

## 2. Results

### 2.1. Count projection is exactly Markov only when aggregate transition rates are fiber-constant

Let *X_t_* denote a time-homogeneous Markov process on a microscopic state space. A state can contain positions, orientations, conformations, contact links, infection states, or species labels. Let

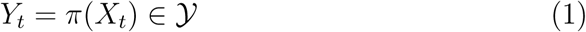

retain only count variables. Equation 1 defines this retained process. For a jump process with microscopic rates *Q*(*x, x*^′^), define the aggregate rate from retained state *y* to *y*^′^ by

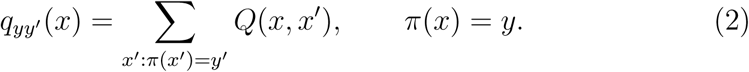

**Theorem 1** (Exact count closure). The projected process Y_t_ is a time-homogeneous Markov chain for every initial distribution if and only if, for every y ≠ *y*^′^*, q_yy_′* (*x*) *is constant over the fiber π*^−1^(*y*)*. The common values define the reduced generator*.

The result is the continuous-time strong-lumpability criterion stated at the level of aggregate count transitions. It permits compensating microscopic channels that produce the same retained transition; equality of every microscopic channel separately is sufficient but not necessary. For a general Markov generator *L*, the equivalent generator condition is that for every suitable function *g* of the retained state, *L*(*g ○ π*) is itself a function of *π*(*x*). A proof and the general generator formulation are given in S1 Text.

Figure 1 makes the fiber-constancy test explicit for a local *A*–*B* interaction. Two spatial configurations *X* and *X*^′^, both belonging to the same fiber *F_y_* and having the same counts (*N_A_, N_B_*) = (2, 2), contain different numbers of reactive contacts. Consequently, their aggregate rates for the same retained transition differ. The same obstruction appears in epidemic and ecological models when count-equivalent states have different susceptible–infectious or predator–prey contact numbers.

**Figure 1:**
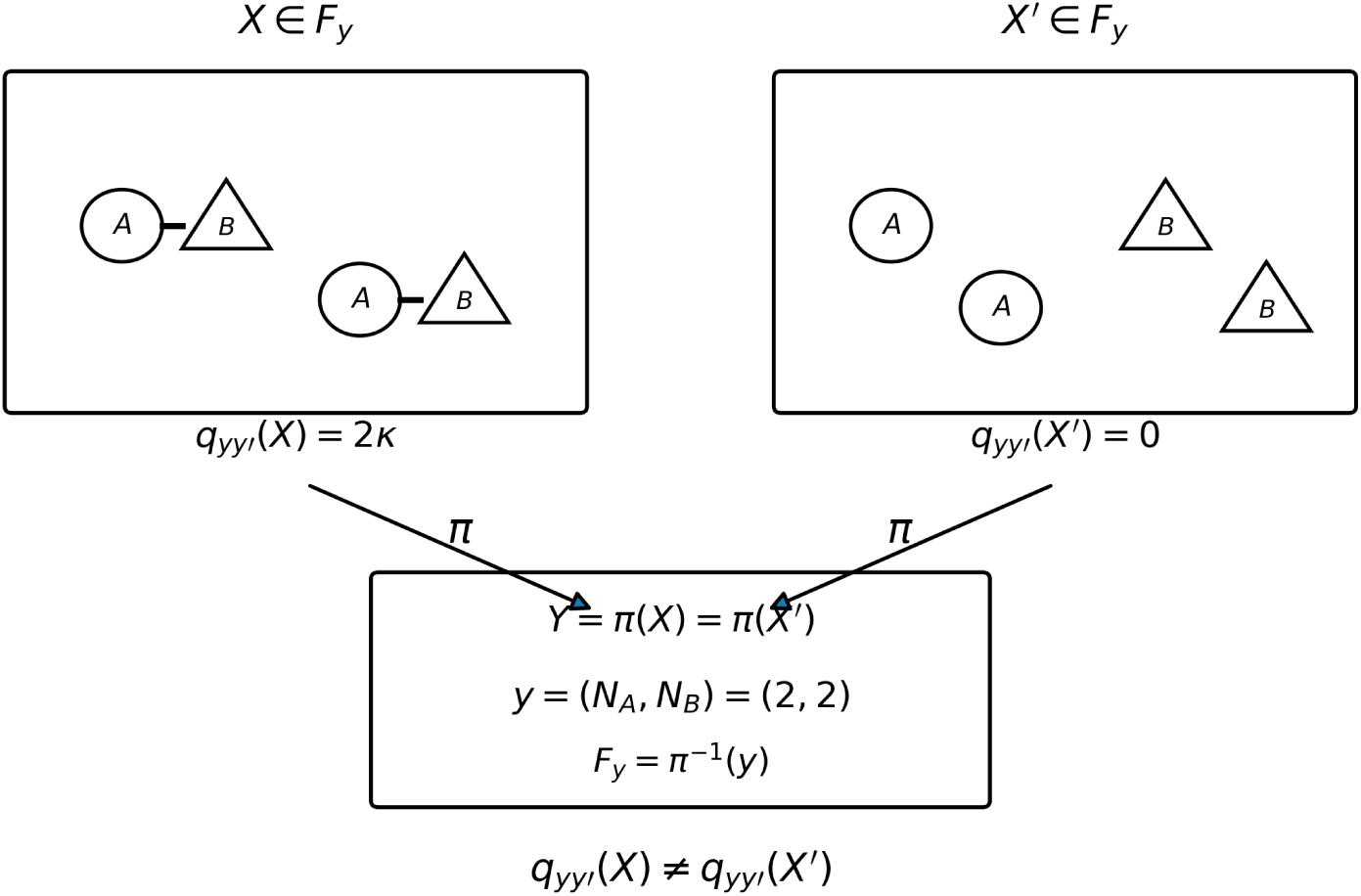
Why identical counts need not define identical dynamics. Two configurations *X, X^′^ F_y_* contain the same numbers of *A* and *B* objects and therefore project to the same zero-dimensional state *y* = (2, 2). In the upper configuration, two local *A*–*B* contacts give aggregate rate 2*κ*; in the lower configuration, spatial separation gives rate zero. Because *q_yy_′* is not constant on the fiber *F_y_*, no exact autonomous count process exists for this transition. Exact count closure requires fiber constancy for every retained transition.

### 2.2. Pair and higher-order correlations locate the missing variables

For pairwise interactions, the evolution of a one-object density generally depends on a two-object density, whose evolution depends on a three-object density. In molecular reaction–diffusion systems this is a variable-particle-number analogue of a BBGKY hierarchy [16, 17, 20]. Analogous hierarchies occur in network epidemics and spatial point-process ecology [5, 18, 19, 21]. For an *A*+*B* interaction with kernel *K*(*z, z*^′^), the mean event rate contains

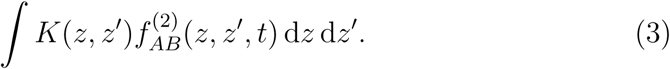

The pair-rate expression in Eq 3 makes the missing correlation explicit. A zero-dimensional mass-action approximation replaces the pair density by a product of one-object densities and then averages *K*. A pair approximation retains *f* ^(2)^ but closes *f* ^(3)^. For a chain 1 − 2 − 3, a common closure is

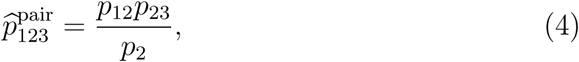

The closure in Eq 4 follows from the probability chain rule plus the approximation 1 ┴3|2; it does not follow from Bayes’ theorem alone. A triplet approximation retains *p*_123_ and may close a four-object chain by 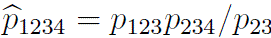.

These levels define a practical hierarchy: deterministic count ODE, stochastic count process, pair closure, triplet closure, and complete spatial process. Bayesian conditioning has a separate role. Given data and count state *y*, the ensemble-dependent effective rate is

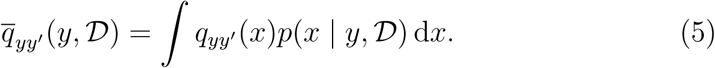

The effective rate in Eq 5 is an optimal conditional mean for a specified ensemble, not an assertion of exact closure.

### 2.3. Information risks distinguish structural non-closure from practical approximation error

Let *µ_y_* be a conditional distribution on a count fiber and 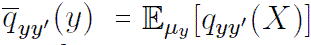. We defined an instantaneous conditional Poisson-rate risk

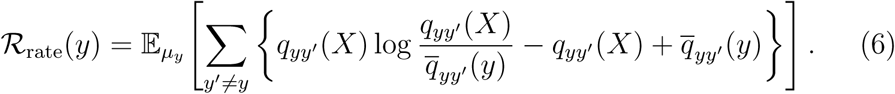

The risk in Eq 6 is nonnegative and is zero precisely when the aggregate rates are constant almost surely under *µ_y_*. If this holds for every distribution on every fiber, the exact condition in Theorem 1 follows.

For a finite observation lag Δ, the predictive risk is

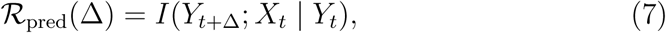

The quantity in Eq 7 measures how much current microscopic information improves prediction beyond the counts. A count-only memory diagnostic is 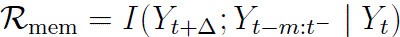. Pair and triplet closure losses can be measured

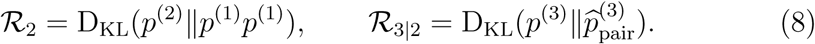

The correlation risks in Eq 8 complement the other diagnostics. The risks answer different questions: *R*_rate_ diagnoses rate heterogeneity on a fiber, *R*_pred_ diagnoses finite-time predictive loss, and *R*_2_ or *R*_3|2_ diagnoses the correlation order omitted by a closure.

### 2.4. An exactly solvable reaction separated exact failure from the fast-mixing limit

We considered one *A* and one *B* molecule in two compartments. The hidden states were colocated (*C*) or separated (*S*). Each molecule changed compartment at rate *d*, while reaction occurred from *C* at rate *k*. Both hidden states projected to counts (1, 1), but their instantaneous reaction rates were *k* and 0, so exact count closure failed.

If *p* is the initial colocation probability, the survival probability was

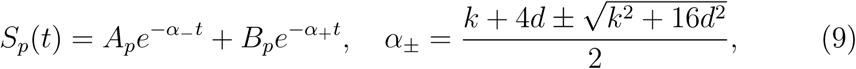

with *A_p_*= (*α*_+_ – *kp*)*/*(*α*_+_ – *α*_−_) and *B_p_* = (*kp* – *α*_−_)*/*(*α*_+_ – *α*_−_). The non-single-exponential survival law demonstrated memory in the count process. As *d/k* → ∞, *α*_−_ → *k/*2 and the fast mode decayed on the mixing time, leaving the well-mixed survival exp(−*kt/*2) (Figs 2, 3, and 4).

**Figure 2:**
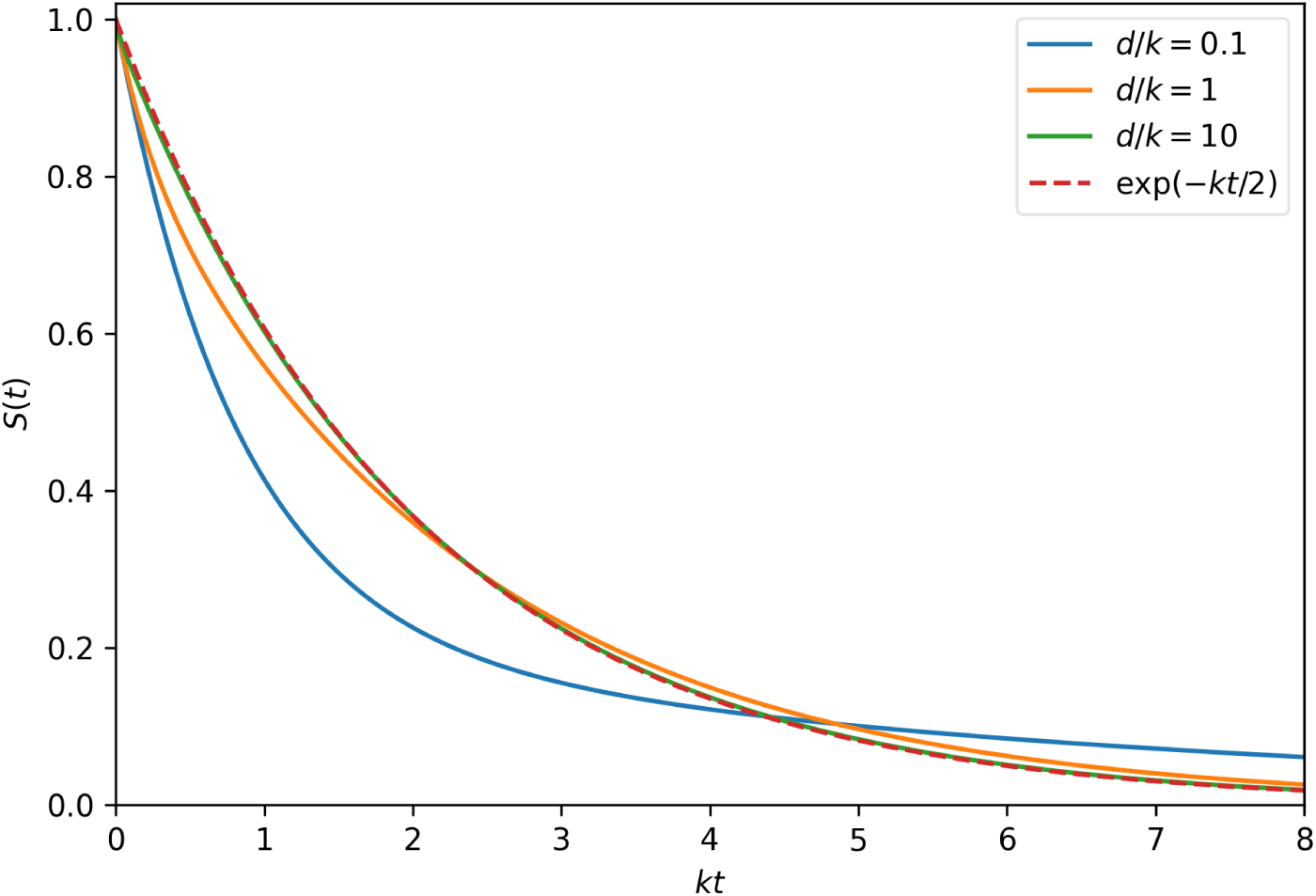
Exact survival in the hidden two-compartment reaction. Slow hidden mixing produced a non-exponential count-level waiting-time law. Fast mixing recovered the effective well-mixed survival.

**Figure 3:**
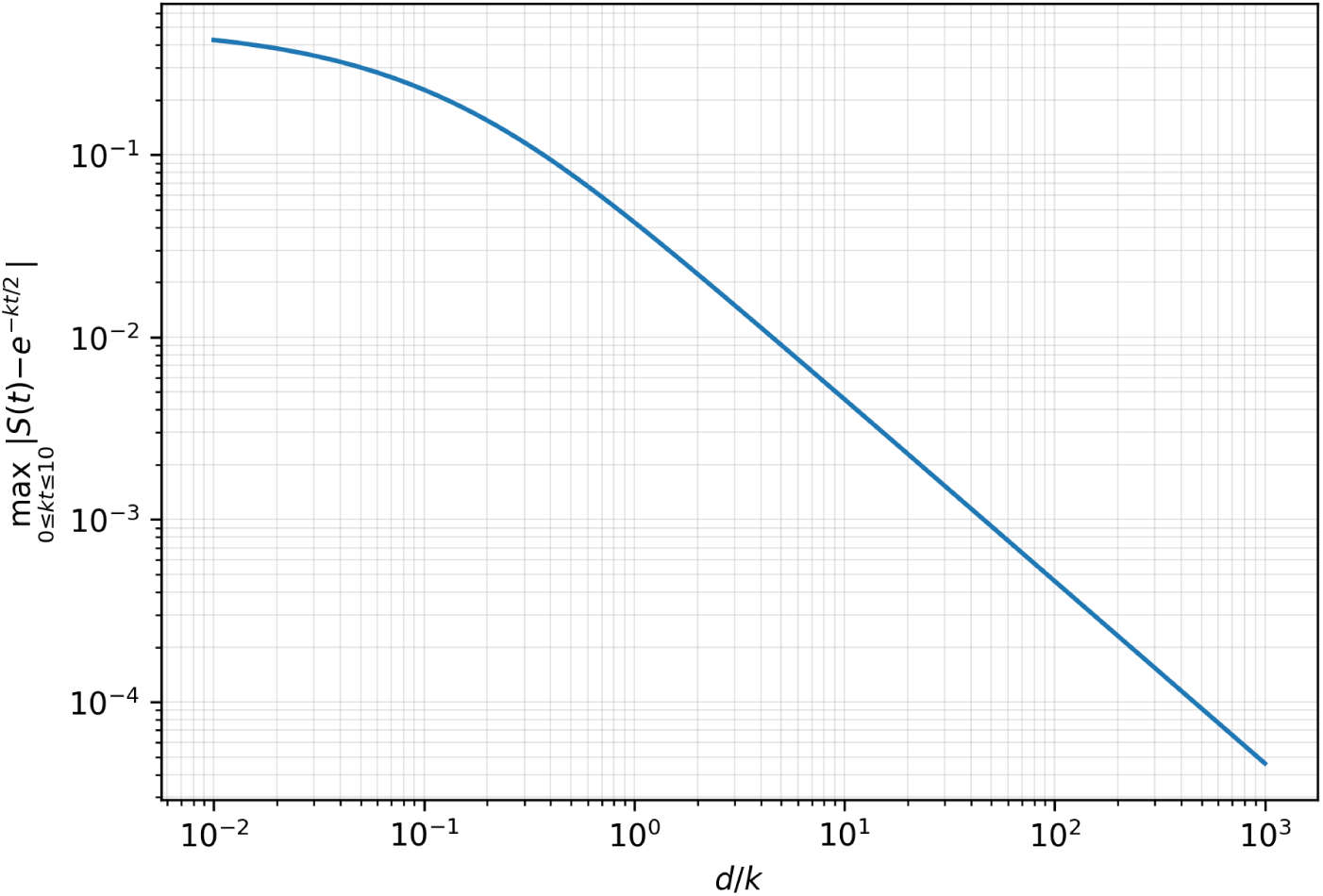
Well-mixed survival error decreased with hidden mixing. The maximum absolute error relative to the fast-mixing law was computed analytically for equilibrium initialization.

**Figure 4:**
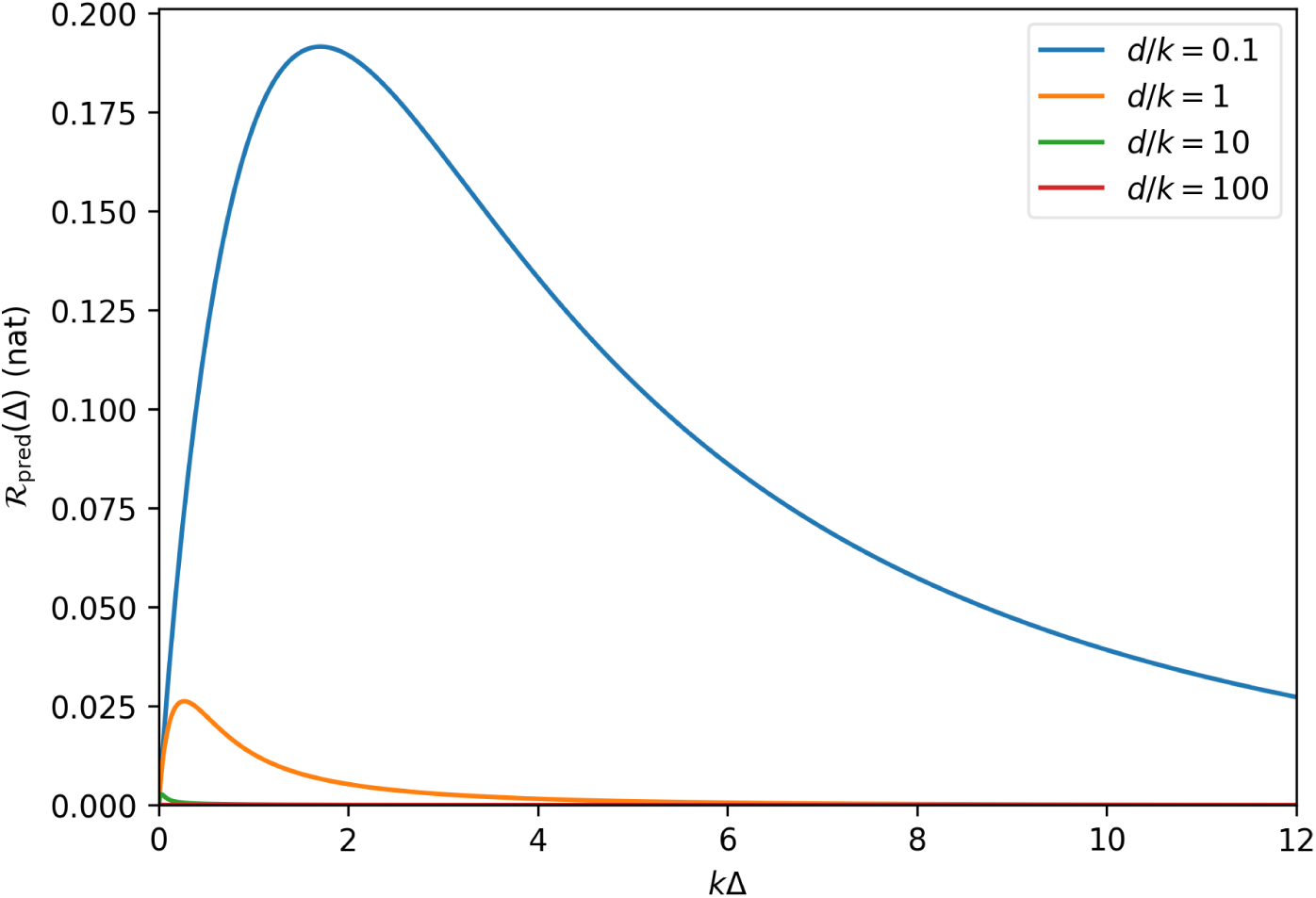
Finite-lag predictive information depended on the mixing ratio. Predictive KL risk was largest when the hidden initial geometry affected reaction over the observation interval and decreased as hidden mixing became fast.

### 2.5. Copy number and mixing imposed distinct constraints on biochemical reductions

For a domain of characteristic size *L* and relative diffusivity *D*_rel_, a first mixing-time estimate is *τ*_mix_ ≃ *L*^2^*/*(*π*^2^*D*_rel_). For a bimolecular reaction with rate constant *k*_2_ and partner concentration *c*, *τ*_int_ ≃ 1*/*(*k*_2_*c*). Their ratio is

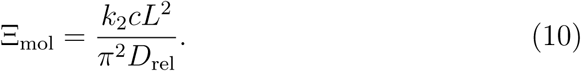

The dimensionless screen in Eq 10 compares interaction and relaxation. Small Ξ_mol_ supports rapid spatial relaxation, but does not guarantee a deterministic ODE. The expected copy number is *N* = *N*_A_*cV*, and intrinsic relative fluctuations are of order *N* ^−1^*^/^*^2^ in a simple Poisson benchmark. Increasing volume therefore reduces copy-number noise while increasing the mixing time as *V* ^2^*^/^*^3^.

Using literature ranges for *Escherichia coli* cell size and cytoplasmic protein diffusion [22–24], we evaluated a representative *V* = 3.2 fL, *L* = 3.0 *µ*m screen. The second-order rate *k*_2_ = 10^6^ M^−1^s^−1^ was used only as a replaceable sensitivity value, not as a measured rate for a particular reaction. The resulting diagram identified a lower concentration imposed by copy number and an upper concentration imposed by spatial mixing (Figs 5 and 6). At sufficiently low diffusivity these bounds crossed, leaving no concentration interval that simultaneously satisfied the selected deterministic-noise and mixing tolerances.

**Figure 5:**
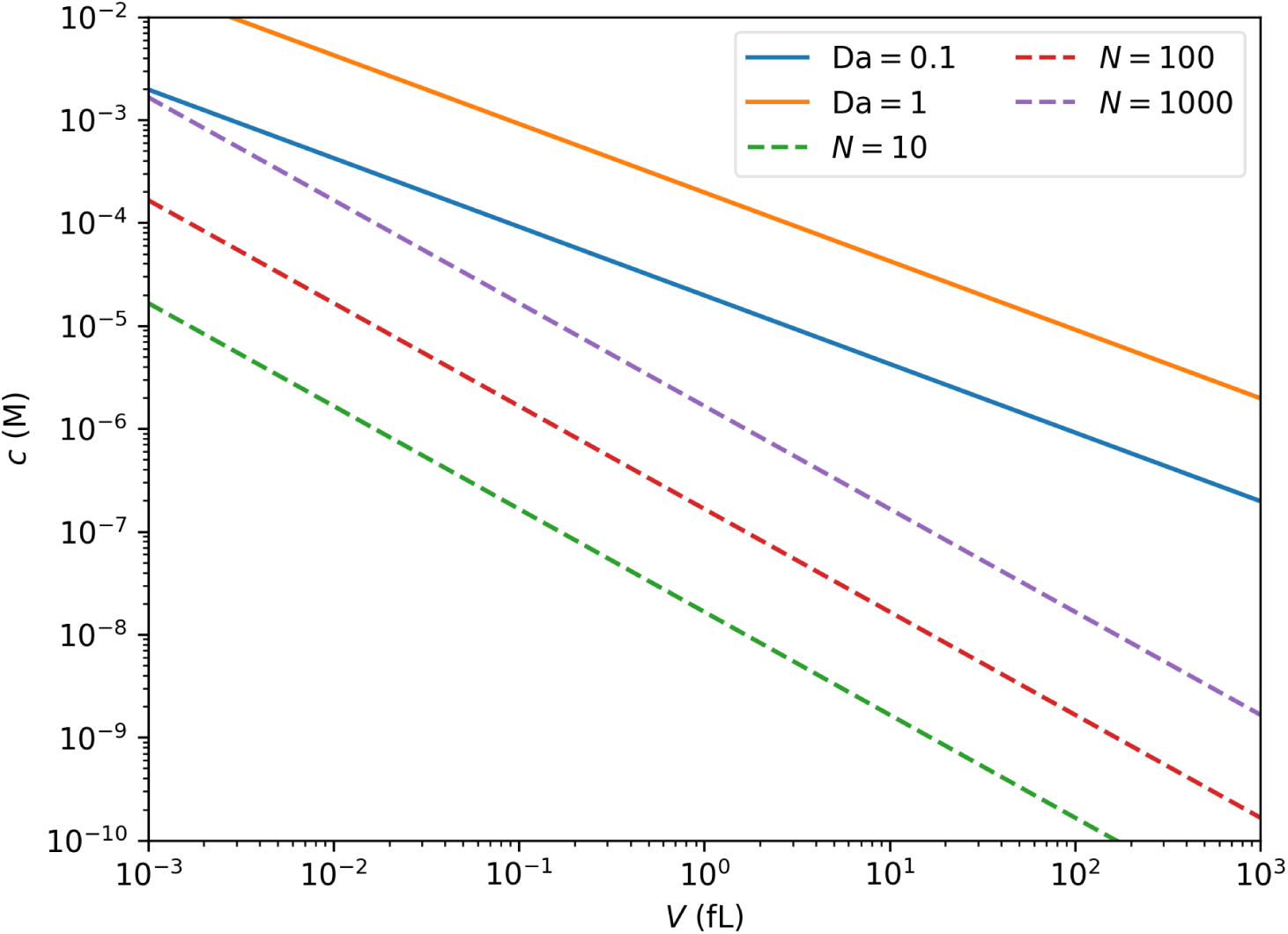
Parameter-level screening separated stochastic and spatial failure modes. Copy number and the mixing–reaction ratio define different constraints; numerical boundaries are tolerance-dependent and are not universal phase transitions.

**Figure 6:**
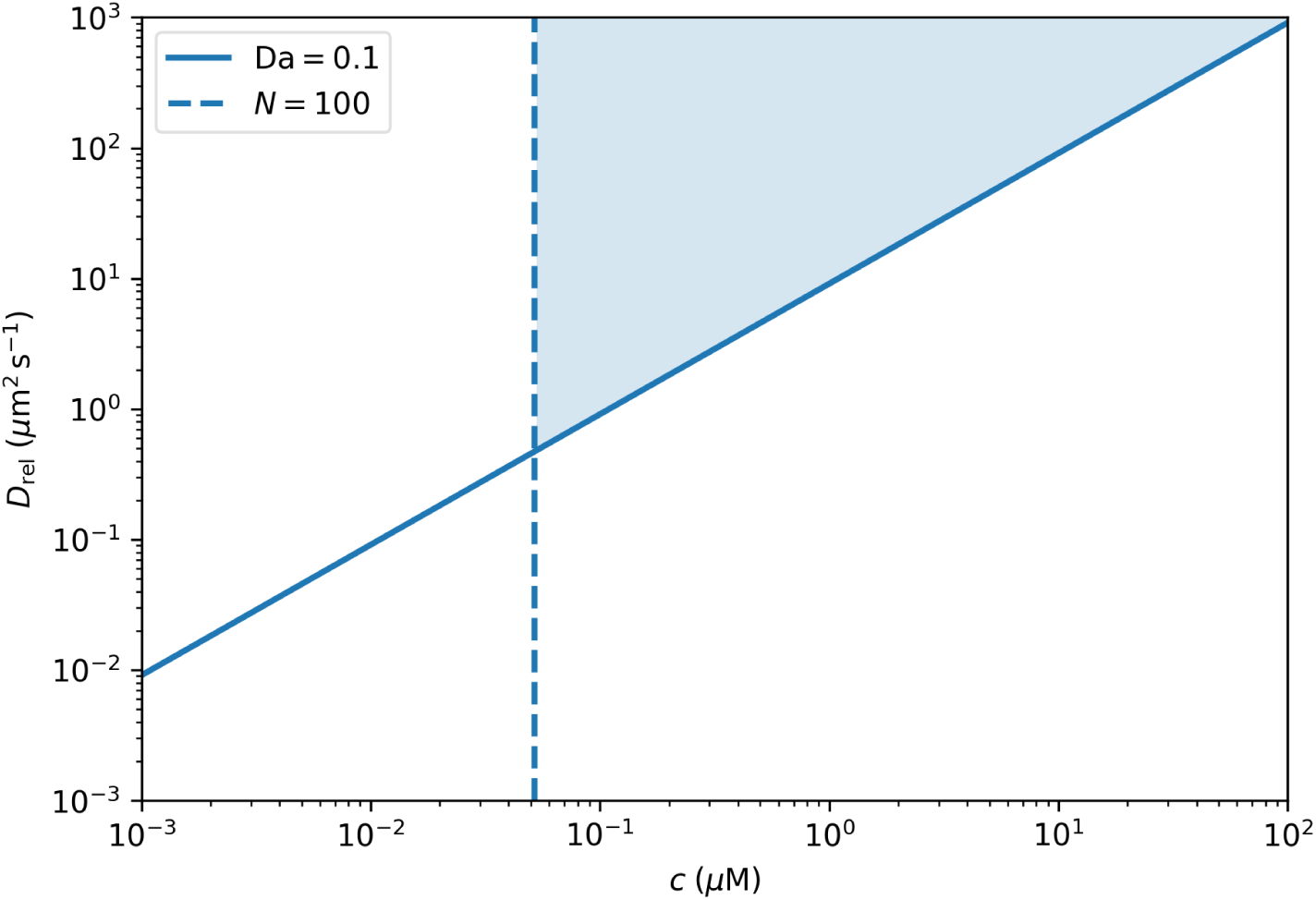
Illustrative *E. coli*-scale deterministic count-model window. The vertical line is the concentration required for 100 expected copies at 3.2 fL. The curve is the relative diffusivity required to keep Ξ_mol_ = 0.1 for the stated sensitivity rate. The shaded region satisfies both selected criteria.

### 2.6. Spatial SIR benchmarks linked contact information to count-model error

We simulated a local SIR process on a periodic two-dimensional lattice. Infection occurred through nearest-neighbour susceptible–infectious contacts, recovery was local-state independent, and a rearrangement process controlled mixing. The retained state was (*S, I, R*). At fixed counts, a clustered infected population and a randomly mixed population had different susceptible–infectious edge counts and therefore different aggregate infection rates.

Across four rearrangement rates and mixed or clustered initial geometry, the early-time pair mutual information between neighboring states ranged from 0.003 to 0.162 nats. The root-mean-square error (RMSE) of the infected fraction relative to the well-mixed SIR ODE ranged from 0.014 to 0.145. Pair risk and RMSE had Pearson correlation 0.90 and Spearman correlation 0.95.

Slow mixing and clustered initialization produced the largest final-size errors, whereas rapid rearrangement reduced both risk and error (Fig 7).

**Figure 7:**
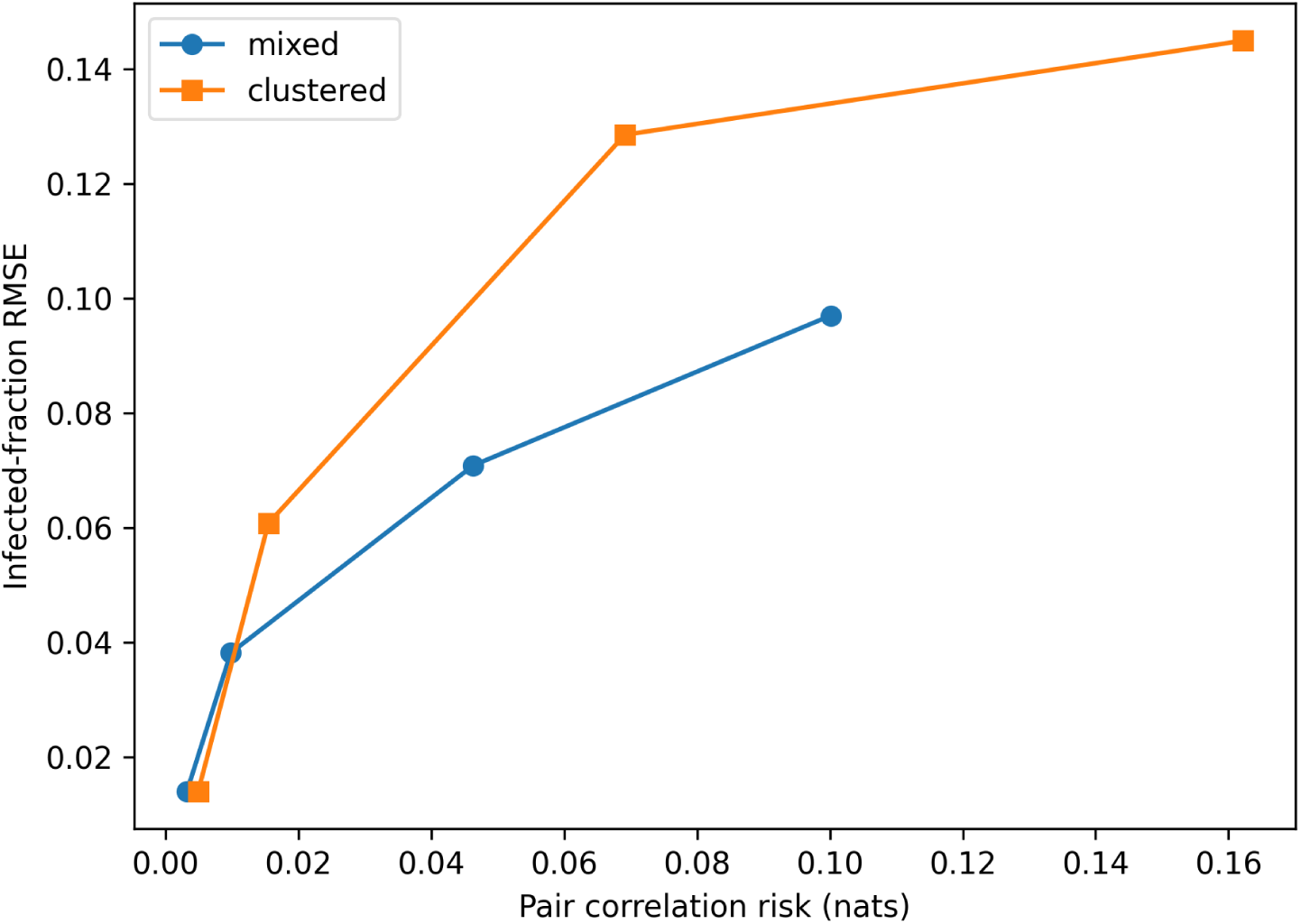
Pair-correlation risk tracked well-mixed SIR error. Each point represents one initial geometry and rearrangement regime. The ordinate is the infected-fraction RMSE between the mean spatial simulation and the deterministic SIR ODE.

### 2.7. Spatial predator–prey benchmarks showed the same closure mechanism

We also simulated a periodic lattice predator–prey contact process with local prey birth, local predation with predator reproduction, predator death, and variable rearrangement. The retained state was the total prey and predator density. Count-equivalent mixed and segregated arrangements generated different predator–prey edge densities and hence different predation rates.

Across the same four rearrangement regimes, pair risk ranged from 0.003 to 0.325 nats, while density RMSE relative to the corresponding mean-field ODE ranged from 0.018 to 0.250. Pair risk and RMSE had Pearson correlation 0.91 and Spearman correlation 1.00. Segregated low-mobility states produced the greatest count-model error (Fig 8). The benchmark is a controlled spatial Lotka–Volterra-type process rather than a calibrated ecosystem model.

**Figure 8:**
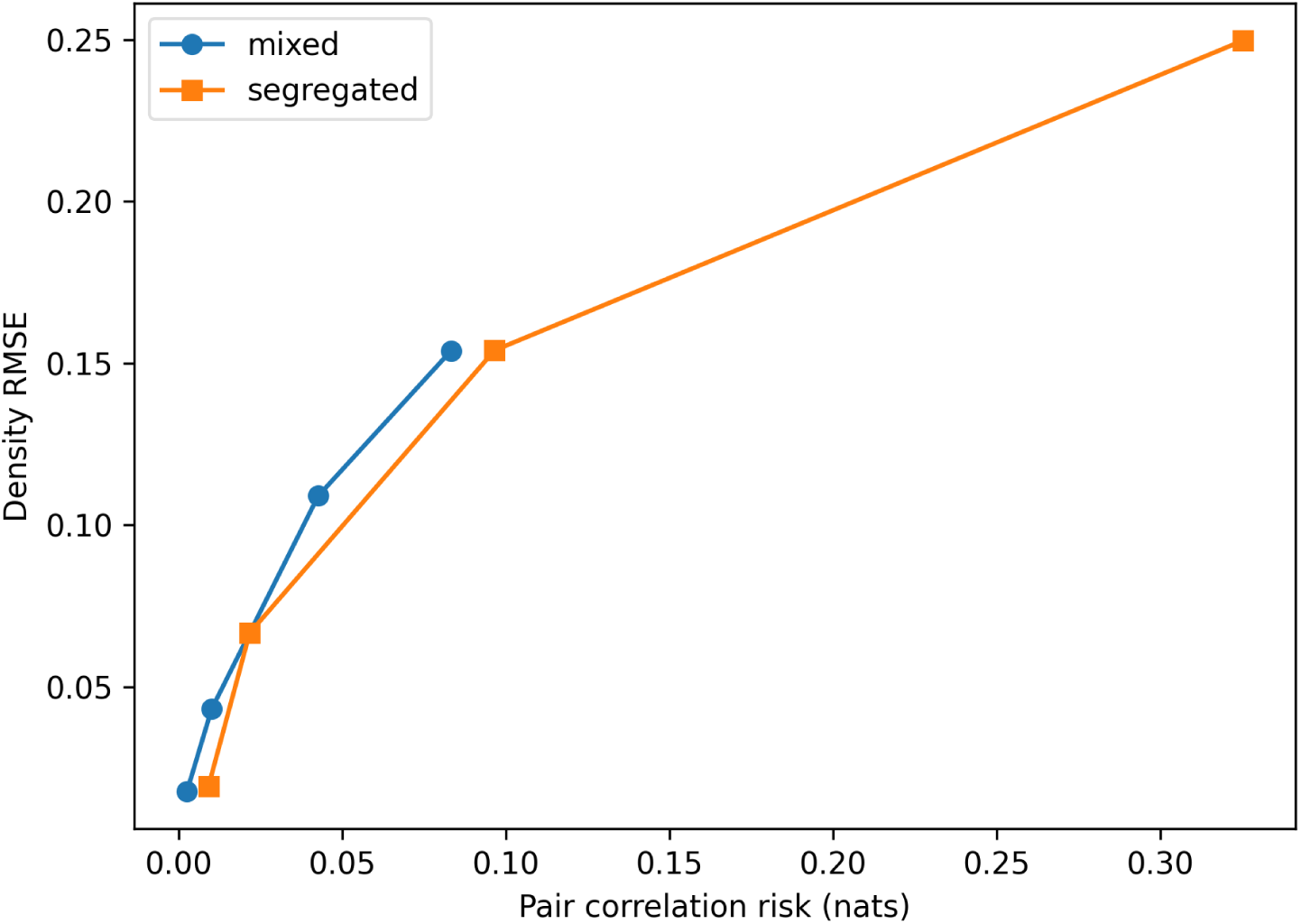
Pair-correlation risk tracked mean-field predator–prey error. Each point represents one initial geometry and rearrangement regime. The error combines prey– and predator-density discrepancies.

### 2.8. Risk–error association transferred across epidemic and ecological benchmarks

After normalizing risk and model error within each benchmark, the pooled Pearson and Spearman correlations were 0.91 and 0.99, respectively (Fig 9). Monte Carlo permutation tests gave *p <* 10^−4^ for both pooled coefficients; bootstrap intervals are reported in S1 Text and the machine-readable robustness table. The normalization prevents the numerical scales of the two models from being interpreted as a universal physical collapse. Rather, the result shows that the same diagnostic principle transferred: when spatial pair information was small, count ODEs were accurate; when pair information was large, count ODE error increased.

**Figure 9:**
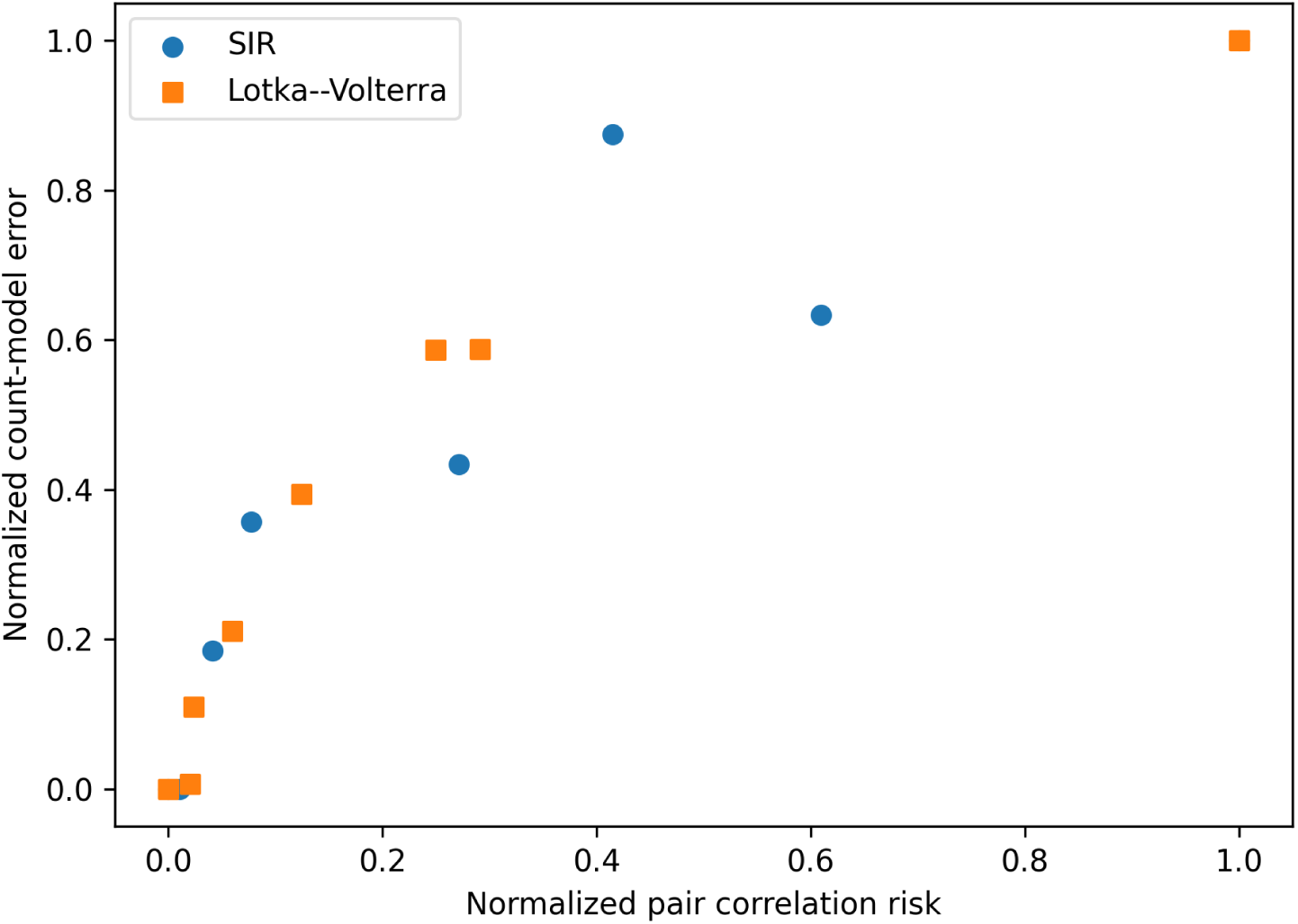
Normalized pair risk and count-model error were associated across two biological scales. Values were normalized separately within each benchmark; the plot demonstrates transfer of ranking, not a universal numerical threshold.

A sensitivity analysis replaced global rearrangement by nearest-neighbour exchange while retaining the local interaction rules. The risk–error ranking remained positive for SIR (Spearman *ρ* = 0.83, permutation *p* = 0.015) and predator–prey dynamics (*ρ* = 0.98, *p <* 10^−3^). These smaller auxiliary simulations do not calibrate real mobility, but they show that the association was not an artifact of globally permuting site labels. Full results and bootstrap intervals are supplied in S1 Text and S1 Data.

## 3. Discussion

Our results support a simple but consequential distinction. The fiber-constancy theorem is the standard strong-lumpability result; the present novelty lies in using it to organize a biological spatial-reduction problem and in connecting its failure to correlation order, information loss, measurable time scales, and cross-domain model error. Large population or molecule number can justify a deterministic approximation to a correctly specified count process, but it cannot by itself justify discarding spatial degrees of freedom. Conversely, fast spatial mixing can support a stochastic count process even when copy numbers are too small for an ODE. The two reductions—spatial process to stochastic counts, then stochastic counts to deterministic ODE—require different conditions.

The exact theorem is deliberately strict: it asks for a Markov count process for every microscopic initialization. Biological modeling often needs only an approximate description for a restricted ensemble and finite observation lag. The KL risks make this weaker question explicit. The rate risk detects heterogeneity that is invisible to counts at infinitesimal resolution. Predictive risk incorporates hidden mixing over a specified lag. Pair and triplet risks identify which correlation order is missing. These measures complement rather than replace conventional error metrics.

The SIR and predator–prey benchmarks strengthened generality because they used the same projection logic outside molecular kinetics. Their purpose was not to propose a new epidemic or ecological model. Instead, they showed that a local-interaction count projection fails for the same reason at distinct scales: total event rates depend on unresolved contact structure. The strong risk–error association is encouraging, but the sample of model classes and parameter regimes remains limited. A universal threshold was neither assumed nor inferred.

The present biochemical example is also intentionally a scale screen rather than a calibrated model of one molecular reaction. A stronger biological validation would use a specific intracellular association with measured copy number, diffusivity, localization, and kinetic parameters, then compare particle, reaction–diffusion master equation, pair, triplet, stochastic count, and ODE predictions. Such validation is the highest-priority next step. Other extensions include weak lumpability for selected initial ensembles, geometry-specific spectral gaps, rotational and conformational mixing, and Bayesian uncertainty in effective closure rates.

Related in-press work by the author considered lossy finite shared states, volatile write–diffuse–decay fields, and memory–responsiveness tradeoffs in unconventional computing [25–27]. Those studies provide conceptual examples of hidden-state compression and environmental memory, but they neither prove nor replace the biological closure results developed here.

The framework suggests a model-selection workflow. First test copy number and mixing–interaction time scales. Second estimate pair information from spatial simulations or imaging data. If pair risk is negligible, a count model may be sufficient. If it is substantial but triplet residual is small, retain pair variables. If higher-order residuals remain substantial, a higher-order or full spatial description is warranted. This workflow turns the statement that “space matters” into a falsifiable and reproducible hierarchy of diagnostics.

## 4. Materials and methods

### 4.1. General spatial process and aggregate-rate criterion

The microscopic state was a typed point or lattice process *X_t_* = (*x_i_, σ_i_, h_i_*), where *x_i_* is position, *σ_i_* is a molecular species, epidemic state, or ecological species, and *h_i_* is an optional hidden internal state. The projection retained counts by type. For each microscopic jump, all channels producing the same count transition were summed before the fiber-constancy test in Eq 2. Full proofs are supplied in S1 Text.

### 4.2. Information-risk estimation

Pair risk was estimated as the mutual information of states across undirected nearest-neighbour lattice edges. For state variables *U* and *V* at the ends of an edge,

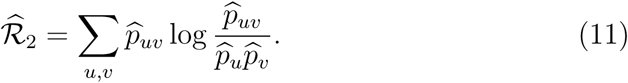

Equation 11 was evaluated from pooled undirected edges. The instantaneous event-rate risk used the Poisson divergence 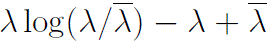, divided by lattice size. Main-text correlations used the pair mutual-information risk because it is common to all local interaction types.

### 4.3. Two-compartment reaction

The hidden-state generator before reaction was

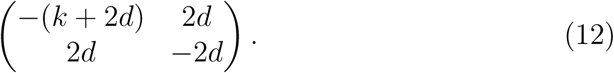

The generator in Eq 12 yields Eq 9 by diagonalization. Scripts evaluated survival error and finite-lag predictive KL risk over mixing ratios.

### 4.4. Spatial SIR benchmark

A 20 × 20 periodic lattice was initialized with 4% infectious sites and the remaining sites susceptible. Initial infectious sites were either uniformly mixed or placed in a compact cluster. Infection of a susceptible site occurred with probability 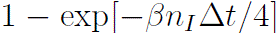, where *n_I_* is the number of infectious nearest neighbours, *β* = 1.5, and Δ*t* = 0.05. Infectious sites recovered with probability 1 exp(-*γ*Δ*t*), with *γ* = 0.4. Rearrangement rates were 0, 0.2, 1, and 5 per unit time; a random subset of sites was permuted at each step to create controlled mixing. Ten replicate simulations were run for each condition. In a supplementary sensitivity analysis, global rearrangement was replaced by stochastic nearest-neighbour exchanges on a smaller lattice, with four replicates per condition. The well-mixed comparator was 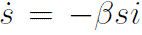, 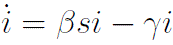 and 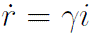, initialized at the ensemble mean.

### 4.5. Spatial predator–prey benchmark

A 24×24 periodic lattice had states empty, prey, or predator. Prey birth into local empty sites had rate *b* = 1.0, local predation with predator reproduction had rate *a* = 2.0, and predator death had rate *d* = 0.55. Mixed and spatially segregated initial states were tested at rearrangement rates 0, 0.2, 1, and 5. Eight replicates were simulated per condition with Δ*t* = 0.05. A supplementary nearest-neighbour exchange analysis used four replicates per condition on a smaller lattice. The mean-field comparator was 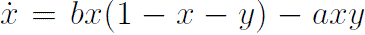 and 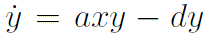, where *x* and *y* are prey and predator densities.

### 4.6. Statistical summaries and reproducibility

Model error was the time-averaged RMSE between ensemble-mean spatial trajectories and their ODE comparators. Pearson and Spearman correlations were calculated across eight conditions in each benchmark. Two-sided Monte Carlo permutation tests used 10,000 label permutations, and percentile bootstrap intervals used 2,000 resamples with fixed seeds. These intervals describe uncertainty across the tested condition set, not population-level sampling uncertainty. All random seeds, scripts, raw trajectory summaries, robustness tables, and plotting code are included in S1 Code and S1 Data. The software uses Python 3 with NumPy and Matplotlib. No human, animal, clinical, or personally identifiable data were used.

## Data availability

All data underlying the findings are provided in machine-readable CSV and JSON files in the accompanying data archive. All author-generated source code required to reproduce the simulations, analyses, and figures is provided in the accompanying code archive under the MIT License. No access restrictions apply.

## CRediT authorship contribution statement

Chikoo Oosawa: Conceptualization, Formal analysis, Investigation, Methodology, Software, Validation, Visualization, Writing–original draft, Writing–review and editing.

## Funding

This work was partially supported by JSPS KAKENHI Grant Number JP26K09271.

## Declaration of competing interest

The author declares that he has no known competing financial interests or personal relationships that could have appeared to influence the work reported in this paper.

## Declaration of generative AI and AI-assisted technologies in the writing process

During the preparation of this work, the author used OpenAI ChatGPT for language editing, structural organization, literature checking, and assistance in drafting reproducibility code. The author independently reviewed the text, verified cited sources, checked mathematical statements and computational outputs, and takes full responsibility for the content.

## Supplementary material

Supplementary Material contains proofs, additional derivations, and limitations of approximate closure diagnostics. Machine-readable data and Python source code are supplied as separate archives.

## Supporting information

Additional theoretical derivations, proofs, model definitions, parameter details, and sensitivity analyses supporting the results/

Python scripts used to reproduce the simulations, numerical analyses, parameter screens, and figures reported in the manuscript.

Machine-readable CSV and JSON data underlying the numerical results and figures.

## Notes

### Competing Interest Statement

The authors have declared no competing interest.

