## Additional theoretical derivations, proofs, model definitions, parameter details, and sensitivity analyses supporting the results/ for "Projection criteria and information risks for zero-dimensional biological dynamics across molecular, epidemic, and ecological systems"

### Supplementary Material: Proofs, derivations, and limitations of information-risk diagnostics

Chikoo Oosawa

#### 1 Strong lumpability for a count projection

Let $X_t$ be a continuous-time Markov chain on a finite or countable state space $\mathcal{X}$ with generator $Q = (Q(x, x'))$, and let $\pi : \mathcal{X} \to \mathcal{Y}$ be a surjective count projection. For $y \neq y'$ and $x \in \pi^{-1}(y)$, define

$$q_{yy'}(x) = \sum_{x':\pi(x')=y'} Q(x, x').$$

**Theorem 1.** *The projected process $Y_t = \pi(X_t)$ is a time-homogeneous Markov chain for every initial distribution on $\mathcal{X}$ if and only if $q_{yy'}(x)$ is constant in $x$ on each fiber $\pi^{-1}(y)$ for every $y \neq y'$. The common values are the off-diagonal entries of the reduced generator.*

*Proof.* Assume fiber constancy and write $\bar{Q}(y, y') = q_{yy'}(x)$ for any $x \in \pi^{-1}(y)$. For any bounded function $g$ on $\mathcal{Y}$,

$$\mathcal{L}(g \circ \pi)(x) = \sum_{x'} Q(x, x')[g(\pi(x')) - g(\pi(x))] = \sum_{y' \neq y} \bar{Q}(y, y')[g(y') - g(y)],$$

where $y = \pi(x)$. The right-hand side depends on $x$ only through $y$, so the subspace of functions constant on fibers is invariant under the generator. The semigroup therefore preserves this subspace and induces the Markov semigroup with generator $\bar{Q}$ on $\mathcal{Y}$.

Conversely, assume $Y_t$ is a homogeneous Markov chain for every initial distribution. Initialize $X_0 = x$ with $\pi(x) = y$. For $y' \neq y$,

$$\Pr(Y_h = y' \mid X_0 = x) = h\, q_{yy'}(x) + o(h).$$

The left-hand side must depend on $x$ only through $y$, because the projected transition law is common to all point-mass initializations in the same fiber. Dividing by $h$ and taking $h \downarrow 0$ gives fiber constancy. □

**Remark 1.** *Equality of every microscopic reaction channel is sufficient but not necessary. Microscopic channels that lead to the same retained transition may compensate; the necessary object is their aggregate rate.*

#### 2 General generator formulation

For a Markov process with generator $\mathcal{L}$, exact Markov closure is equivalent, under the usual domain and uniqueness assumptions, to invariance of the algebra of retained observables:

$$\mathcal{L}(g \circ \pi) = \bar{\mathcal{L}} g \circ \pi$$

for all suitable $g$. This formulation includes diffusions and hybrid jump–diffusion processes. It also clarifies why count observables fail to close when the generator produces pair densities, local contact counts, orientation-dependent kernels, or other functions that are not determined by the retained counts.

#### 3 Conditional Poisson-rate risk

Let $\mu_y$ be a conditional ensemble on a fiber and let $q_j(X)$ denote the aggregate rates of retained transitions. Define $\bar{q}_j = \mathbb{E}_{\mu_y} q_j(X)$. The relative entropy rate between independent Poisson clocks with intensities $q_j(X)$ and $\bar{q}_j$ is

$$\mathcal{R}_{\mathrm{rate}}(y) = \mathbb{E}_{\mu_y} \sum_j \left[ q_j(X) \log \frac{q_j(X)}{\bar{q}_j} - q_j(X) + \bar{q}_j \right].$$

For $a, b \geq 0$, the scalar divergence $a \log(a/b) - a + b$ is nonnegative, with equality exactly at $a = b$ (with the usual conventions at zero). Hence $\mathcal{R}_{\mathrm{rate}} \geq 0$. It vanishes if and only if every retained transition rate is constant $\mu_y$-almost surely. Requiring this for every probability measure supported on every fiber recovers the pointwise strong-lumpability criterion.

#### 4 Finite-time prediction and memory

The predictive risk

$$\mathcal{R}_{\mathrm{pred}}(\Delta) = I(Y_{t+\Delta}; X_t \mid Y_t)$$

is zero exactly when the microscopic state gives no additional one-step predictive information at lag $\Delta$ once the count state is known. Vanishing for a single lag and a single ensemble is weaker than strong lumpability. The count-only memory risk

$$\mathcal{R}_{\mathrm{mem}}(\Delta, m) = I(Y_{t+\Delta}; Y_{t-m:t^-} \mid Y_t)$$

can be estimated without observing the microscopic state. A positive value detects non-Markovianity of the observed count process, but it does not identify which hidden variable is responsible.

#### 5 Correlation hierarchy and closure risks

For pairwise interactions, a one-object density generally depends on a two-object density, and the latter on a three-object density. A mean-field approximation replaces $p^{(2)}$ by $p^{(1)}p^{(1)}$. A chain pair closure replaces

$$p_{123} \quad \text{by} \quad \widehat{p}_{123} = \frac{p_{12}p_{23}}{p_2},$$

which is equivalent to the additional approximation $1 \perp 3 \mid 2$. Bayes' theorem organizes conditional distributions and posterior updating, but does not by itself justify this conditional-independence statement.

Natural residual risks are

$$\mathcal{R}_2 = D_{\mathrm{KL}}(p^{(2)} \| p^{(1)}p^{(1)}), \qquad \mathcal{R}_{3|2} = D_{\mathrm{KL}}(p^{(3)} \| \widehat{p}^{(3)}_{\mathrm{pair}}).$$

Small $\mathcal{R}_2$ supports a count or mean-field description for the measured observable; large $\mathcal{R}_2$ with small $\mathcal{R}_{3|2}$ supports retaining pairs. These are diagnostics rather than universal error bounds. Their relation to a chosen prediction error depends on the interaction kernel, observation horizon, state occupancy, and sampling accuracy.

#### 6 Two-compartment reaction derivation

Let $C$ and $S$ denote hidden colocated and separated states. Before absorption by reaction, the transient generator is

$$A = \begin{pmatrix} -(k+2d) & 2d \\ 2d & -2d \end{pmatrix}.$$

The positive decay rates are the roots of

$$\alpha^2 - (k+4d)\alpha + 2kd = 0,$$

namely

$$\alpha_\pm = \frac{k + 4d \pm \sqrt{k^2 + 16d^2}}{2}.$$

If $p$ is the initial probability of $C$, the survival law has the form

$$S_p(t) = A_p e^{-\alpha_- t} + B_p e^{-\alpha_+ t},$$

where $A_p = (\alpha_+ - kp)/(\alpha_+ - \alpha_-)$ and $B_p = (kp - \alpha_-)/(\alpha_+ - \alpha_-)$. Unless coefficients or rates degenerate, this is not a single exponential and therefore cannot be generated by an autonomous one-state count process with a constant reaction hazard. For $d/k \to \infty$, $\alpha_- \to k/2$ while $\alpha_+ \sim 4d$, so after the fast boundary layer the effective survival is $e^{-kt/2}$.

#### 7 Parameter screens

For a domain with slowest nonconstant diffusion mode $\lambda_1$, a mixing estimate is $\tau_{\text{mix}} \simeq 1/(D_{\text{rel}}\lambda_1)$. The main text uses the cube-like estimate $\lambda_1 \simeq \pi^2/L^2$, giving

$$\Xi_{\text{mol}} = \frac{k_2 c L^2}{\pi^2 D_{\text{rel}}}.$$

This is a screening ratio, not a universal transition point. Geometry, reflecting or absorbing boundaries, anomalous diffusion, reversible binding, membrane localization, and orientation or conformational relaxation can alter the relevant relaxation time. Deterministic ODE validity additionally requires adequate copy number and dynamical stability against noise-driven switching or extinction.

#### 8 Benchmark interpretation

The SIR and predator–prey benchmarks were designed to isolate one mechanism: local pair interactions create count-equivalent states with different transition rates. Random site permutation is used as a controlled rearrangement operator. It is not intended as a literal model of human movement or organismal dispersal. A sensitivity analysis replaced global rearrangement by nearest-neighbour exchanges. Pair-risk and ODE-error rankings remained positively associated in both model classes, indicating that the main qualitative result was not specific to global permutation. The observed correlations establish transfer of the diagnostic across controlled models, not a universal calibration for real epidemics or ecosystems.

#### 9 Finite-condition robustness analysis

For each main benchmark, uncertainty in the association across the eight tested conditions was assessed by two-sided Monte Carlo permutation tests with 10,000 label permutations and percentile bootstrap intervals with 2,000 resamples. The resulting intervals are descriptive for the finite condition set; they do not imply that the conditions were sampled from a biological population. For global rearrangement, the SIR Spearman coefficient was 0.952 (permutation $p = 0.0015$; bootstrap 95% interval $[0.620, 1.000]$), and the predator–prey coefficient was 1.000 ($p < 10^{-4}$; interval $[1.000, 1.000]$). After within-system normalization, the pooled coefficient was 0.987 ($p < 10^{-4}$; interval $[0.925, 1.000]$).

Under nearest-neighbour exchange, the corresponding coefficients were 0.833 for SIR ($p = 0.0146$; interval $[0.308, 1.000]$) and 0.976 for predator–prey dynamics ($p = 0.0004$; interval $[0.730, 1.000]$). These auxiliary calculations used smaller systems and fewer replicates than the main benchmarks and are therefore interpreted only as a mechanism-sensitivity check. Machine-readable results are provided in the following files:

`robustness_statistics.csv`
`sir_local_exchange_summary.csv`
`lv_local_exchange_summary.csv`.

#### 10 Limitations and recommended validation

The principal limitations are: (i) no specific intracellular reaction with a complete experimentally constrained parameter set; (ii) no theorem converting pair KL risk to a universal trajectory-error bound; (iii) finite lattices and a small number of parameter regimes; (iv) empirical pair mutual information can depend on binning, neighborhood definition, and sampling; and (v) higher-order closure risks were defined but not benchmarked numerically. A decisive biological validation should compare particle or reaction–diffusion simulations, stochastic count models, pair and triplet closures, and ODEs for a named system with measured copy numbers, localization, diffusion, and reaction kinetics.
