## Supplementary material for "Projection criteria and information risks for zero-dimensional biological dynamics across molecular, epidemic, and ecological systems": Python scripts used to reproduce the simulations, numerical analyses, parameter screens, and figures reported in the manuscript.: REPRODUCIBILITY.md

# Reproducibility package

This package accompanies the manuscript:

**Projection criteria and information risks for zero-dimensional biological dynamics across molecular, epidemic, and ecological systems**

## Contents

- `manuscript_R7_MB.tex` and `.pdf`: review-format journal manuscript.
- `manuscript_R7_bioRxiv.tex` and `.pdf`: reader-oriented preprint build with equivalent scientific content.
- `supplement/Supplementary_Material.tex` and `.pdf`: proofs, derivations, robustness results, and limitations.
- `scripts/`: exact two-compartment calculations, parameter screening, spatial SIR, spatial predator–prey, local-exchange sensitivity, robustness statistics, and plotting scripts.
- `data/`: machine-readable CSV and JSON outputs.
- `figures/`: all main-text figures in PDF and PNG formats.
- `S1_Code.zip` and `S1_Data.zip`: upload-ready supporting archives.

## Environment

Python 3.10 or later is recommended. Install dependencies with:

```bash
python3 -m venv .venv
source .venv/bin/activate
python3 -m pip install -r requirements.txt
```

## Reproduce results

Run the wrapper from the package root:

```bash
python3 scripts/run_all.py
```

The wrapper launches each component in an isolated Python process. Runtime is machine dependent. All random-number seeds are fixed. If a system-specific library or process issue occurs, run the components independently in the order below:

```bash
python3 scripts/local_exchange_sensitivity.py
python3 scripts/spatial_sir_benchmark.py
python3 scripts/spatial_lv_benchmark.py
python3 scripts/two_compartment_model.py --out data/two_compartment_summary.csv
python3 scripts/generate_fig1_projection.py
python3 scripts/generate_fig2_survival.py
python3 scripts/generate_fig3_error.py
python3 scripts/generate_fig4_predictive_kl.py
python3 scripts/generate_fig5_parameter_map.py
python3 scripts/generate_fig6_ecoli_window.py
python3 scripts/generate_cross_scale_figures.py
python3 scripts/robustness_statistics.py
```

## Compile documents

```bash
latexmk -pdf manuscript_R7_MB.tex
latexmk -pdf manuscript_R7_bioRxiv.tex
cd supplement
latexmk -pdf Supplementary_Material.tex
```

## Scope of the benchmarks

The SIR and predator–prey calculations are controlled methodological tests, not forecasts of a particular outbreak or ecosystem. Global permutation provides a transparent mixing control; nearest-neighbour exchange is included as a sensitivity analysis. The Escherichia coli calculation is a parameter screen; its second-order rate constant is a replaceable sensitivity value rather than a measurement for a specific intracellular reaction. Bootstrap intervals describe variation over the finite tested condition set, not population-level sampling uncertainty.

## License

Author-generated code is released under the MIT License.
