## Supplementary material for "Projection criteria and information risks for zero-dimensional biological dynamics across molecular, epidemic, and ecological systems": Python scripts used to reproduce the simulations, numerical analyses, parameter screens, and figures reported in the manuscript.: ecoli_0d_risk.py

```
#!/usr/bin/env python3
"""Screen zero-dimensional validity for an E. coli-scale compartment.

All thresholds are diagnostics, not universal error guarantees.
"""
from __future__ import annotations
import argparse, json, math

NA = 6.02214076e23

def screen(volume_fl: float, length_um: float, concentration_uM: float,
           drel_um2_s: float, k2_M_inv_s: float = 1e6,
           delta: float = 0.1, eps_sp: float = 0.1) -> dict[str, float | bool]:
    c_M = concentration_uM * 1e-6
    v_L = volume_fl * 1e-15
    n = NA * c_M * v_L
    rel_noise = 1.0 / math.sqrt(n) if n > 0 else math.inf
    da = k2_M_inv_s * c_M * length_um**2 / (math.pi**2 * drel_um2_s)
    chi_copy = 1.0 / (n * delta**2) if n > 0 else math.inf
    chi_space = da / eps_sp
    return {
        "volume_fL": volume_fl,
        "length_um": length_um,
        "concentration_uM": concentration_uM,
        "Drel_um2_s": drel_um2_s,
        "k2_M_inv_s": k2_M_inv_s,
        "copy_number": n,
        "poisson_relative_noise": rel_noise,
        "spatial_Damkohler": da,
        "chi_copy": chi_copy,
        "chi_space": chi_space,
        "chi_0D": max(chi_copy, chi_space),
        "passes_selected_thresholds": max(chi_copy, chi_space) <= 1.0,
    }

def main() -> None:
    ap = argparse.ArgumentParser()
    ap.add_argument('--volume-fl', type=float, default=3.2)
    ap.add_argument('--length-um', type=float, default=3.0)
    ap.add_argument('--concentration-uM', type=float, default=1.0)
    ap.add_argument('--drel', type=float, default=10.0, help='relative diffusivity in um^2/s')
    ap.add_argument('--k2', type=float, default=1e6, help='M^-1 s^-1')
    ap.add_argument('--delta', type=float, default=0.1)
    ap.add_argument('--eps-sp', type=float, default=0.1)
    args = ap.parse_args()
    print(json.dumps(screen(args.volume_fl, args.length_um, args.concentration_uM,
                            args.drel, args.k2, args.delta, args.eps_sp), indent=2))
if __name__ == '__main__':
    main()
```
