## Supplementary material for "Projection criteria and information risks for zero-dimensional biological dynamics across molecular, epidemic, and ecological systems": Python scripts used to reproduce the simulations, numerical analyses, parameter screens, and figures reported in the manuscript.: generate_cross_scale_figures.py

```
#!/usr/bin/env python3
from pathlib import Path
import csv
import numpy as np
import matplotlib.pyplot as plt
ROOT=Path(__file__).resolve().parents[1]; DATA=ROOT/'data'; FIG=ROOT/'figures';FIG.mkdir(exist_ok=True)

def read(name):
    with (DATA/name).open() as f:return list(csv.DictReader(f))

def scatter(rows,xkey,ykey,out,xlab,ylab,modekey):
    fig,ax=plt.subplots(figsize=(6.2,4.5))
    groups={}
    for r in rows:groups.setdefault(r[modekey],[]).append(r)
    marks=['o','s','^','D']
    for (g,rr),mk in zip(groups.items(),marks):
      x=np.array([float(z[xkey]) for z in rr]);y=np.array([float(z[ykey]) for z in rr])
      idx=np.argsort(x);ax.plot(x[idx],y[idx],marker=mk,label=g)
    ax.set_xlabel(xlab);ax.set_ylabel(ylab);ax.legend(frameon=True,framealpha=0.6)
    fig.tight_layout();fig.savefig(FIG/(out+'.png'),dpi=300);fig.savefig(FIG/(out+'.pdf'));plt.close(fig)

sir=read('sir_benchmark_summary.csv');lv=read('lv_benchmark_summary.csv')
scatter(sir,'pair_KL_nats','infected_RMSE','fig7_sir_risk_error',r'Pair correlation risk (nats)',r'Infected-fraction RMSE','initial_geometry')
scatter(lv,'pair_KL_nats','density_RMSE','fig8_lv_risk_error',r'Pair correlation risk (nats)',r'Density RMSE','initial_geometry')
# pooled normalized association
pts=[]
for system,rows,xk,yk in [('SIR',sir,'pair_KL_nats','infected_RMSE'),('Lotka--Volterra',lv,'pair_KL_nats','density_RMSE')]:
    xs=np.array([float(r[xk]) for r in rows]);ys=np.array([float(r[yk]) for r in rows])
    xn=(xs-xs.min())/(xs.max()-xs.min()+1e-12);yn=(ys-ys.min())/(ys.max()-ys.min()+1e-12)
    pts.extend([(system,x,y) for x,y in zip(xn,yn)])
fig,ax=plt.subplots(figsize=(6.2,4.5))
for sys,mk in [('SIR','o'),('Lotka--Volterra','s')]:
    q=[p for p in pts if p[0]==sys];ax.scatter([p[1] for p in q],[p[2] for p in q],marker=mk,label=sys)
ax.set_xlabel(r'Normalized pair correlation risk');ax.set_ylabel(r'Normalized count-model error');ax.legend(frameon=True,framealpha=0.6)
fig.tight_layout();fig.savefig(FIG/'fig9_cross_scale.png',dpi=300);fig.savefig(FIG/'fig9_cross_scale.pdf');plt.close(fig)
with (DATA/'cross_scale_points.csv').open('w',newline='') as f:
    w=csv.writer(f);w.writerow(['system','normalized_pair_risk','normalized_count_model_error']);w.writerows(pts)
```
