## Supplementary material for "Projection criteria and information risks for zero-dimensional biological dynamics across molecular, epidemic, and ecological systems": Python scripts used to reproduce the simulations, numerical analyses, parameter screens, and figures reported in the manuscript.: generate_fig1_projection.py

```
#!/usr/bin/env python3
"""Generate Fig. 1: count-equivalent microstates with unequal aggregate rates."""
from pathlib import Path
import matplotlib.pyplot as plt
from matplotlib.patches import FancyBboxPatch, FancyArrowPatch, Circle, RegularPolygon

ROOT = Path(__file__).resolve().parents[1]
out = ROOT / 'figures'
out.mkdir(exist_ok=True)
fig, ax = plt.subplots(figsize=(7.4, 5.25))
ax.set_xlim(0, 8)
ax.set_ylim(0.7, 7)
ax.axis('off')

def rounded(x, y, w, h):
    patch = FancyBboxPatch((x, y), w, h, boxstyle='round,pad=0.04',
                           fill=False, linewidth=1.4)
    ax.add_patch(patch)

def draw_a(x, y):
    ax.add_patch(Circle((x, y), 0.22, facecolor='white', edgecolor='black', linewidth=1.3))
    ax.text(x, y, r'$A$', ha='center', va='center', fontsize=10)

def draw_b(x, y):
    ax.add_patch(RegularPolygon((x, y), numVertices=3, radius=0.28,
                                orientation=0, facecolor='white', edgecolor='black', linewidth=1.3))
    ax.text(x, y-0.018, r'$B$', ha='center', va='center', fontsize=9)

rounded(0.35, 4.25, 3.1, 2.0)
rounded(4.55, 4.25, 3.1, 2.0)
ax.text(1.90, 6.52, r'$X\in F_y$', ha='center', fontsize=14)
ax.text(6.10, 6.52, r"$X'\in F_y$", ha='center', fontsize=14)

draw_a(0.95, 5.35); draw_b(1.52, 5.35)
draw_a(2.25, 4.90); draw_b(2.82, 4.90)
ax.plot([1.17, 1.28], [5.35, 5.35], color='black', lw=2.2)
ax.plot([2.47, 2.58], [4.90, 4.90], color='black', lw=2.2)
draw_a(5.15, 5.35); draw_a(5.75, 4.85)
draw_b(6.65, 5.35); draw_b(7.20, 4.85)

ax.text(1.90, 4.00, r'$q_{yy\prime}(X)=2\kappa$', ha='center', fontsize=13)
ax.text(6.10, 4.00, r"$q_{yy\prime}(X')=0$", ha='center', fontsize=13)

ax.add_patch(FancyArrowPatch((1.95, 3.78), (3.55, 3.00), arrowstyle='-|>',
                             mutation_scale=15, linewidth=1.35))
ax.add_patch(FancyArrowPatch((6.05, 3.78), (4.45, 3.00), arrowstyle='-|>',
                             mutation_scale=15, linewidth=1.35))
ax.text(2.80, 3.42, r'$\pi$', ha='center', fontsize=15)
ax.text(5.20, 3.42, r'$\pi$', ha='center', fontsize=15)

rounded(2.35, 1.85, 3.3, 1.35)
ax.text(4.00, 2.88, r"$Y=\pi(X)=\pi(X')$", ha='center', fontsize=13)
ax.text(4.00, 2.42, r'$y=(N_A,N_B)=(2,2)$', ha='center', fontsize=13)
ax.text(4.00, 2.02, r'$F_y=\pi^{-1}(y)$', ha='center', fontsize=12)
ax.text(4.00, 1.36, r"$q_{yy\prime}(X)\ne q_{yy\prime}(X')$", ha='center', fontsize=14)

fig.tight_layout(pad=0.25)
fig.savefig(out/'fig1_projection.png', dpi=300, bbox_inches='tight')
fig.savefig(out/'fig1_projection.pdf', bbox_inches='tight')
plt.close(fig)
```
