## Supplementary material for "Projection criteria and information risks for zero-dimensional biological dynamics across molecular, epidemic, and ecological systems": Python scripts used to reproduce the simulations, numerical analyses, parameter screens, and figures reported in the manuscript.: generate_fig2_survival.py

```
#!/usr/bin/env python3
from pathlib import Path
import sys
import numpy as np
import matplotlib.pyplot as plt
sys.path.insert(0, str(Path(__file__).resolve().parent))
from two_compartment_model import survival

ROOT = Path(__file__).resolve().parents[1]
OUT = ROOT / "figures"
DATA = ROOT / "data"
OUT.mkdir(exist_ok=True); DATA.mkdir(exist_ok=True)

k = 1.0
x = np.linspace(0.0, 8.0, 1201)
ratios = [0.1, 1.0, 10.0]
fig, ax = plt.subplots(figsize=(6.3,4.4))
cols = [x]
header = ["kt"]
for ratio in ratios:
    s = survival(x/k, k, ratio*k, 1.0)
    ax.plot(x, s, label=rf"$d/k={ratio:g}$")
    cols.append(s); header.append(f"S_d_over_k_{ratio:g}")
ref = np.exp(-0.5*x)
ax.plot(x, ref, linestyle="--", label=r"$\exp(-kt/2)$")
cols.append(ref); header.append("well_mixed_reference")
ax.set_xlabel(r"$kt$")
ax.set_ylabel(r"$S(t)$")
ax.set_xlim(0,8); ax.set_ylim(0,1.02)
ax.legend(frameon=True, framealpha=0.5)
fig.tight_layout()
for ext in ("pdf","png"):
    fig.savefig(OUT / f"fig2_survival.{ext}", dpi=300, bbox_inches="tight")
np.savetxt(DATA / "fig2_survival_data.csv", np.column_stack(cols), delimiter=",", header=",".join(header), comments="")
plt.close(fig)
```
