## Supplementary material for "Projection criteria and information risks for zero-dimensional biological dynamics across molecular, epidemic, and ecological systems": Python scripts used to reproduce the simulations, numerical analyses, parameter screens, and figures reported in the manuscript.: generate_fig3_error.py

```
#!/usr/bin/env python3
from pathlib import Path
import sys
import numpy as np
import matplotlib.pyplot as plt
sys.path.insert(0, str(Path(__file__).resolve().parent))
from two_compartment_model import survival

ROOT = Path(__file__).resolve().parents[1]
OUT = ROOT / "figures"
DATA = ROOT / "data"
OUT.mkdir(exist_ok=True); DATA.mkdir(exist_ok=True)

k = 1.0
ratios = np.logspace(-2,3,180)
t = np.linspace(0.0,10.0,5001)
ref = np.exp(-0.5*t)
errs = []
for ratio in ratios:
    s = survival(t, k, ratio*k, 0.5)
    errs.append(np.max(np.abs(s-ref)))
errs = np.asarray(errs)
fig, ax = plt.subplots(figsize=(6.3,4.4))
ax.loglog(ratios, errs)
ax.set_xlabel(r"$d/k$")
ax.set_ylabel(r"$\max_{0\leq kt\leq 10}|S(t)-e^{-kt/2}|$")
ax.grid(True, which="both", linewidth=0.4, alpha=0.4)
fig.tight_layout()
for ext in ("pdf","png"):
    fig.savefig(OUT / f"fig3_error.{ext}", dpi=300, bbox_inches="tight")
np.savetxt(DATA / "fig3_error_data.csv", np.column_stack([ratios,errs]), delimiter=",", header="d_over_k,max_abs_error", comments="")
plt.close(fig)
```
