## Supplementary material for "Projection criteria and information risks for zero-dimensional biological dynamics across molecular, epidemic, and ecological systems": Python scripts used to reproduce the simulations, numerical analyses, parameter screens, and figures reported in the manuscript.: generate_fig4_predictive_kl.py

```
#!/usr/bin/env python3
from pathlib import Path
import sys
import numpy as np
import matplotlib.pyplot as plt

HERE = Path(__file__).resolve().parent
ROOT = HERE.parent
sys.path.insert(0, str(HERE))
from kl_risk import predictive_kl

OUT = ROOT / "figures"
DATA = ROOT / "data"
OUT.mkdir(exist_ok=True)
DATA.mkdir(exist_ok=True)

k = 1.0
x = np.linspace(0.0, 12.0, 1601)
ratios = [0.1, 1.0, 10.0, 100.0]
cols = [x]
header = ["k_delta"]

fig, ax = plt.subplots(figsize=(6.3, 4.4))
for ratio in ratios:
    risk = predictive_kl(x / k, k, ratio * k, 0.5)
    ax.plot(x, risk, label=rf"$d/k={ratio:g}$")
    cols.append(risk)
    header.append(f"Rpred_d_over_k_{ratio:g}")
ax.set_xlabel(r"$k\Delta$")
ax.set_ylabel(r"$\mathcal{R}_{\mathrm{pred}}(\Delta)$ (nat)")
ax.set_xlim(0.0, 12.0)
ax.set_ylim(bottom=0.0)
ax.legend(frameon=True, framealpha=0.5)
fig.tight_layout()
for ext in ("pdf", "png"):
    fig.savefig(OUT / f"fig4_predictive_kl.{ext}", dpi=300, bbox_inches="tight")
np.savetxt(DATA / "fig4_predictive_kl_data.csv", np.column_stack(cols), delimiter=",", header=",".join(header), comments="")
plt.close(fig)
```
