## Supplementary material for "Projection criteria and information risks for zero-dimensional biological dynamics across molecular, epidemic, and ecological systems": Python scripts used to reproduce the simulations, numerical analyses, parameter screens, and figures reported in the manuscript.: generate_fig5_parameter_map.py

```
#!/usr/bin/env python3
from pathlib import Path
import numpy as np
import matplotlib.pyplot as plt

ROOT = Path(__file__).resolve().parents[1]
OUT = ROOT / "figures"
DATA = ROOT / "data"
OUT.mkdir(exist_ok=True)
DATA.mkdir(exist_ok=True)

# Illustrative bimolecular parameters, explicitly stated in the manuscript caption.
k2 = 1.0e6       # M^-1 s^-1
Drel = 20.0      # micrometre^2 s^-1
NA = 6.02214076e23

V = np.logspace(-3, 3, 800)  # fL = micrometre^3
L2 = V ** (2.0 / 3.0)

fig, ax = plt.subplots(figsize=(6.3, 4.7))
for da, ls in [(0.1, "-"), (1.0, "-")]:
    c = da * np.pi**2 * Drel / (k2 * L2)
    ax.loglog(V, c, linestyle=ls, label=rf"$\mathrm{{Da}}={da:g}$")
for n, ls in [(10.0, "--"), (100.0, "--"), (1000.0, "--")]:
    # c[M] * V[fL] * 1e-15 L/fL * NA = N
    c = n / (NA * V * 1.0e-15)
    ax.loglog(V, c, linestyle=ls, label=rf"$N={n:g}$")
ax.set_xlabel(r"$V$ (fL)")
ax.set_ylabel(r"$c$ (M)")
ax.set_xlim(1.0e-3, 1.0e3)
ax.set_ylim(1.0e-10, 1.0e-2)
ax.legend(frameon=True, framealpha=0.5, ncol=2)
fig.tight_layout()
for ext in ("pdf", "png"):
    fig.savefig(OUT / f"fig5_parameter_map.{ext}", dpi=300, bbox_inches="tight")

rows = []
for v in np.logspace(-3, 3, 61):
    for c in np.logspace(-10, -2, 81):
        n = NA * c * v * 1.0e-15
        da = k2 * c * v**(2.0/3.0) / (np.pi**2 * Drel)
        rows.append((v, c, n, da))
np.savetxt(DATA / "fig5_parameter_map_data.csv", np.asarray(rows), delimiter=",", header="V_fL,c_M,N,Da", comments="")
plt.close(fig)
```
