## Supplementary material for "Projection criteria and information risks for zero-dimensional biological dynamics across molecular, epidemic, and ecological systems": Python scripts used to reproduce the simulations, numerical analyses, parameter screens, and figures reported in the manuscript.: generate_fig6_ecoli_window.py

```
#!/usr/bin/env python3
from pathlib import Path
import csv, math
import numpy as np
import matplotlib.pyplot as plt

ROOT = Path(__file__).resolve().parents[1]
FIG = ROOT / 'figures'
DATA = ROOT / 'data'
FIG.mkdir(exist_ok=True); DATA.mkdir(exist_ok=True)

V = 3.2
L = 3.0
k2 = 1e6
delta = 0.1
eps = 0.1
NA = 6.02214076e23
c_noise = delta**-2 / (NA * V * 1e-15) * 1e6  # uM
c = np.logspace(-3, 2, 500)
d_boundary = k2 * (c*1e-6) * L**2 / (eps * math.pi**2)

fig, ax = plt.subplots(figsize=(6.4,4.5))
ax.set_xscale('log'); ax.set_yscale('log')
ax.plot(c, d_boundary, linewidth=2, label=r'$\mathrm{Da}=0.1$')
ax.axvline(c_noise, linewidth=2, linestyle='--', label=r'$N=100$')
ax.fill_between(c, d_boundary, 1e3, where=(c>=c_noise), alpha=0.18)
ax.set_xlim(1e-3,1e2); ax.set_ylim(1e-3,1e3)
ax.set_xlabel(r'$c\;(\mu\mathrm{M})$')
ax.set_ylabel(r'$D_{\mathrm{rel}}\;(\mu\mathrm{m}^2\,\mathrm{s}^{-1})$')
ax.legend(frameon=True, framealpha=0.6, loc='upper left')
fig.tight_layout()
fig.savefig(FIG/'fig6_ecoli_window.pdf')
fig.savefig(FIG/'fig6_ecoli_window.png', dpi=300)
plt.close(fig)

with (DATA/'fig6_ecoli_window_data.csv').open('w', newline='') as f:
    w=csv.writer(f); w.writerow(['concentration_uM','Drel_boundary_um2_s','copy_threshold_uM'])
    for ci,di in zip(c,d_boundary): w.writerow([ci,di,c_noise])

rows=[]
for ci in [0.01,0.1,1.0,10.0]:
    N=NA*ci*1e-6*V*1e-15
    rows.append([ci,N,N**-0.5, k2*ci*1e-6*L**2/(math.pi**2*10), k2*ci*1e-6*L**2/(math.pi**2*1)])
with (DATA/'ecoli_case_summary.csv').open('w', newline='') as f:
    w=csv.writer(f); w.writerow(['concentration_uM','copy_number','relative_noise','Da_Drel10','Da_Drel1']); w.writerows(rows)
```
