## Supplementary material for "Projection criteria and information risks for zero-dimensional biological dynamics across molecular, epidemic, and ecological systems": Python scripts used to reproduce the simulations, numerical analyses, parameter screens, and figures reported in the manuscript.: kl_risk.py

```
#!/usr/bin/env python3
"""Information-risk diagnostics for the exact two-compartment benchmark.

The hidden unreacted state is C (molecules colocated) or S (separated).
Reaction occurs only from C at rate k, while C <-> S mixing occurs at rate 2d.
The count projection sees only whether reaction has occurred.
"""
from __future__ import annotations

import numpy as np
from two_compartment_model import survival


def binary_entropy(p: np.ndarray | float) -> np.ndarray:
    p = np.asarray(p, dtype=float)
    out = np.zeros_like(p)
    mask = (p > 0.0) & (p < 1.0)
    out[mask] = -p[mask] * np.log(p[mask]) - (1.0 - p[mask]) * np.log(1.0 - p[mask])
    return out


def predictive_kl(delta: np.ndarray, k: float, d: float, p_colocated: float = 0.5) -> np.ndarray:
    """I(Y_delta; X_0 | Y_0=unreacted), in nats.

    Y_delta is binary (unreacted/reacted); X_0 is C or S.
    """
    delta = np.asarray(delta, dtype=float)
    if k <= 0.0 or d < 0.0:
        raise ValueError("Require k>0 and d>=0")
    if not 0.0 <= p_colocated <= 1.0:
        raise ValueError("p_colocated must be in [0,1]")
    r_c = 1.0 - survival(delta, k, d, 1.0)
    r_s = 1.0 - survival(delta, k, d, 0.0)
    r_bar = p_colocated * r_c + (1.0 - p_colocated) * r_s
    return binary_entropy(r_bar) - p_colocated * binary_entropy(r_c) - (1.0 - p_colocated) * binary_entropy(r_s)


def instantaneous_rate_kl(k: float, p_colocated: np.ndarray | float) -> np.ndarray:
    """Conditional Poisson-rate KL risk k p log(1/p), in nats/time."""
    p = np.asarray(p_colocated, dtype=float)
    if k <= 0.0 or np.any((p < 0.0) | (p > 1.0)):
        raise ValueError("Require k>0 and p in [0,1]")
    out = np.zeros_like(p)
    mask = p > 0.0
    out[mask] = k * p[mask] * np.log(1.0 / p[mask])
    return out
```
