## Supplementary material for "Projection criteria and information risks for zero-dimensional biological dynamics across molecular, epidemic, and ecological systems": Python scripts used to reproduce the simulations, numerical analyses, parameter screens, and figures reported in the manuscript.: local_exchange_sensitivity.py

```
#!/usr/bin/env python3
"""Nearest-neighbour exchange sensitivity for SIR and predator-prey benchmarks.

The main benchmarks use controlled global rearrangement. This script replaces
that operation by local nearest-neighbour swaps while preserving all other
model definitions, providing a robustness check against the mixing mechanism.
"""
from __future__ import annotations
from pathlib import Path
import csv, math
import numpy as np
from scipy.stats import pearsonr, spearmanr

ROOT = Path(__file__).resolve().parents[1]
DATA = ROOT / 'data'
DATA.mkdir(exist_ok=True)


def neighbours(a):
    return (np.roll(a,1,0), np.roll(a,-1,0), np.roll(a,1,1), np.roll(a,-1,1))


def pair_mi(state, n_states=3):
    pairs=[]
    for b in (np.roll(state,-1,0), np.roll(state,-1,1)):
        pairs.append(np.stack([state.ravel(), b.ravel()], axis=1))
    p=np.concatenate(pairs, axis=0)
    joint=np.zeros((n_states,n_states), float)
    for i,j in p:
        joint[int(i),int(j)] += 1
    joint /= joint.sum()
    pi=joint.sum(axis=1); pj=joint.sum(axis=0)
    out=0.0
    for i in range(n_states):
        for j in range(n_states):
            if joint[i,j] > 0 and pi[i] > 0 and pj[j] > 0:
                out += joint[i,j]*math.log(joint[i,j]/(pi[i]*pj[j]))
    return out


def local_swaps(state, mobility, dt, rng):
    n_events = int(rng.poisson(mobility * state.size * dt))
    L0, L1 = state.shape
    for _ in range(n_events):
        i = int(rng.integers(L0)); j = int(rng.integers(L1)); d = int(rng.integers(4))
        if d == 0: ni, nj = (i+1) % L0, j
        elif d == 1: ni, nj = (i-1) % L0, j
        elif d == 2: ni, nj = i, (j+1) % L1
        else: ni, nj = i, (j-1) % L1
        state[i,j], state[ni,nj] = state[ni,nj], state[i,j]


def sir_init(L, frac_i, mode, rng):
    n=L*L; ni=max(1,int(round(frac_i*n))); s=np.zeros((L,L),np.int8)
    if mode=='mixed': s.ravel()[rng.choice(n,ni,replace=False)] = 1
    else:
        cx=cy=L//2
        coords=sorted(((i,j) for i in range(L) for j in range(L)),
                      key=lambda z:(z[0]-cx)**2+(z[1]-cy)**2)
        for i,j in coords[:ni]: s[i,j]=1
    return s


def sir_sim(L,beta,gamma,mobility,mode,T,dt,seed):
    rng=np.random.default_rng(seed); s=sir_init(L,0.04,mode,rng)
    steps=int(round(T/dt)); save=2; tr=[]; risks=[]; times=[]
    for step in range(steps+1):
        if step%save==0:
            times.append(step*dt); tr.append([np.mean(s==0),np.mean(s==1),np.mean(s==2)])
            risks.append(pair_mi(s))
        if step==steps: break
        local_swaps(s,mobility,dt,rng)
        inf=sum((x==1).astype(int) for x in neighbours(s))
        infect=(s==0)&(rng.random(s.shape)<(1-np.exp(-beta*inf*dt/4)))
        recover=(s==1)&(rng.random(s.shape)<(1-math.exp(-gamma*dt)))
        s[recover]=2; s[infect]=1
    return np.array(times),np.array(tr),np.array(risks)


def sir_ode(beta,gamma,y0,times):
    y=np.array(y0,float); out=[y.copy()]
    def f(v):
        s,i,r=v; return np.array([-beta*s*i,beta*s*i-gamma*i,gamma*i])
    for a,b in zip(times[:-1],times[1:]):
        h=b-a; k1=f(y); k2=f(y+h*k1/2); k3=f(y+h*k2/2); k4=f(y+h*k3)
        y=y+h*(k1+2*k2+2*k3+k4)/6; y=np.maximum(y,0); y/=y.sum(); out.append(y.copy())
    return np.array(out)


def lv_init(L,mode,rng):
    s=np.zeros((L,L),np.int8)
    if mode=='mixed':
        u=rng.random((L,L)); s[u<0.45]=1; s[(u>=0.45)&(u<0.60)]=2
    else:
        s[:,:L//2]=1; mask=rng.random((L,L//2))<0.30; s[:,L//2:][mask]=2
        s[rng.random((L,L))<0.15]=0
    return s


def lv_sim(L,b,a,d,mobility,mode,T,dt,seed):
    rng=np.random.default_rng(seed); s=lv_init(L,mode,rng)
    steps=int(round(T/dt)); save=4; tr=[]; risks=[]; times=[]
    for step in range(steps+1):
        if step%save==0:
            times.append(step*dt); tr.append([np.mean(s==0),np.mean(s==1),np.mean(s==2)])
            risks.append(pair_mi(s))
        if step==steps: break
        local_swaps(s,mobility,dt,rng)
        prey=sum((x==1).astype(int) for x in neighbours(s))
        pred=sum((x==2).astype(int) for x in neighbours(s))
        birth=(s==0)&(rng.random(s.shape)<(1-np.exp(-b*prey*dt/4)))
        eaten=(s==1)&(rng.random(s.shape)<(1-np.exp(-a*pred*dt/4)))
        death=(s==2)&(rng.random(s.shape)<(1-math.exp(-d*dt)))
        s[death]=0; s[birth]=1; s[eaten]=2
    return np.array(times),np.array(tr),np.array(risks)


def lv_ode(b,a,d,y0,times):
    y=np.array(y0,float); out=[y.copy()]
    def f(v):
        e,x,p=v; birth=b*x*e; pred=a*x*p; death=d*p
        return np.array([death-birth,birth-pred,pred-death])
    for t0,t1 in zip(times[:-1],times[1:]):
        h=t1-t0; k1=f(y); k2=f(y+h*k1/2); k3=f(y+h*k2/2); k4=f(y+h*k3)
        y=y+h*(k1+2*k2+2*k3+k4)/6; y=np.maximum(y,0); y/=max(y.sum(),1e-12); out.append(y.copy())
    return np.array(out)


def main():
    mobilities=[0.0,0.2,1.0,5.0]
    sir_rows=[]
    for mode in ['mixed','clustered']:
        for m in mobilities:
            dat=[]; risks=[]
            for r in range(4):
                t,tr,ri=sir_sim(16,1.5,0.4,m,mode,12.0,0.05,61000+r+int(100*m)+(0 if mode=='mixed' else 5000))
                dat.append(tr); risks.append(ri)
            mean=np.mean(dat,0); od=sir_ode(1.5,0.4,mean[0],t)
            risk=float(np.mean(np.mean(risks,0)[:max(2,len(t)//3)]))
            rmse=float(np.sqrt(np.mean((mean[:,1]-od[:,1])**2)))
            sir_rows.append([mode,m,risk,rmse])
    with (DATA/'sir_local_exchange_summary.csv').open('w',newline='') as f:
        w=csv.writer(f); w.writerow(['initial_geometry','mobility','pair_KL_nats','infected_RMSE']); w.writerows(sir_rows)

    lv_rows=[]
    for mode in ['mixed','segregated']:
        for m in mobilities:
            dat=[]; risks=[]
            for r in range(4):
                t,tr,ri=lv_sim(18,1.0,2.0,0.55,m,mode,15.0,0.05,71000+r+int(100*m)+(0 if mode=='mixed' else 5000))
                dat.append(tr); risks.append(ri)
            mean=np.mean(dat,0); od=lv_ode(1.0,2.0,0.55,mean[0],t)
            risk=float(np.mean(np.mean(risks,0)[:max(2,len(t)//3)]))
            rmse=float(np.sqrt(np.mean((mean[:,1:]-od[:,1:])**2)))
            lv_rows.append([mode,m,risk,rmse])
    with (DATA/'lv_local_exchange_summary.csv').open('w',newline='') as f:
        w=csv.writer(f); w.writerow(['initial_geometry','mobility','pair_KL_nats','density_RMSE']); w.writerows(lv_rows)

    out=[]
    for name,rows in [('SIR',sir_rows),('Lotka--Volterra',lv_rows)]:
        x=np.array([r[2] for r in rows]); y=np.array([r[3] for r in rows])
        pr=pearsonr(x,y); sr=spearmanr(x,y)
        out.append([name,float(pr.statistic),float(pr.pvalue),float(sr.statistic),float(sr.pvalue)])
    with (DATA/'local_exchange_correlations.csv').open('w',newline='') as f:
        w=csv.writer(f); w.writerow(['system','pearson_r','pearson_p','spearman_rho','spearman_p']); w.writerows(out)
    print(DATA/'local_exchange_correlations.csv')

if __name__=='__main__': main()
```
