## Supplementary material for "Projection criteria and information risks for zero-dimensional biological dynamics across molecular, epidemic, and ecological systems": Python scripts used to reproduce the simulations, numerical analyses, parameter screens, and figures reported in the manuscript.: robustness_statistics.py

```
#!/usr/bin/env python3
"""Monte Carlo permutation tests and bootstrap confidence intervals."""
from __future__ import annotations
from pathlib import Path
import csv, json
import numpy as np
from scipy.stats import rankdata
ROOT=Path(__file__).resolve().parents[1]; DATA=ROOT/'data'; DATA.mkdir(exist_ok=True)

def read_csv(name,xkey,ykey):
    with (DATA/name).open() as f: rows=list(csv.DictReader(f))
    return np.array([float(r[xkey]) for r in rows]),np.array([float(r[ykey]) for r in rows])

def corr(a,b):
    a=np.asarray(a,float); b=np.asarray(b,float); a=a-a.mean(); b=b-b.mean()
    den=np.sqrt(np.dot(a,a)*np.dot(b,b)); return float(np.dot(a,b)/den) if den>0 else np.nan

def transform(a,kind): return rankdata(a) if kind=='spearman' else np.asarray(a,float)

def mc_perm(x,y,kind,B=10000):
    x=transform(x,kind); y=transform(y,kind); obs=corr(x,y); rng=np.random.default_rng(20260806)
    vals=np.empty(B)
    for i in range(B): vals[i]=corr(x,rng.permutation(y))
    p=(1+np.sum(np.abs(vals)>=abs(obs)-1e-15))/(B+1)
    return obs,float(p)

def boot(x,y,kind,B=2000):
    rng=np.random.default_rng(20260807); vals=[]; n=len(x)
    for _ in range(B):
        idx=rng.integers(0,n,n); a=transform(x[idx],kind); b=transform(y[idx],kind); v=corr(a,b)
        if np.isfinite(v): vals.append(v)
    return [float(z) for z in np.percentile(vals,[2.5,97.5])]

def summarize(label,x,y):
    pr,pp=mc_perm(x,y,'pearson'); sr,sp=mc_perm(x,y,'spearman')
    return {'system':label,'n':len(x),'pearson_r':pr,'pearson_perm_p':pp,'pearson_bootstrap_95':boot(x,y,'pearson'),
            'spearman_rho':sr,'spearman_perm_p':sp,'spearman_bootstrap_95':boot(x,y,'spearman')}

def main():
    inputs=[('SIR-global-rearrangement','sir_benchmark_summary.csv','pair_KL_nats','infected_RMSE'),('Lotka-Volterra-global-rearrangement','lv_benchmark_summary.csv','pair_KL_nats','density_RMSE'),('Pooled-normalized','cross_scale_points.csv','normalized_pair_risk','normalized_count_model_error'),('SIR-local-exchange','sir_local_exchange_summary.csv','pair_KL_nats','infected_RMSE'),('Lotka-Volterra-local-exchange','lv_local_exchange_summary.csv','pair_KL_nats','density_RMSE')]
    res=[]
    for label,name,xk,yk in inputs:
        if (DATA/name).exists(): res.append(summarize(label,*read_csv(name,xk,yk)))
    (DATA/'robustness_statistics.json').write_text(json.dumps(res,indent=2),encoding='utf-8')
    with (DATA/'robustness_statistics.csv').open('w',newline='') as f:
        w=csv.writer(f); w.writerow(['system','n','pearson_r','pearson_perm_p','pearson_ci_low','pearson_ci_high','spearman_rho','spearman_perm_p','spearman_ci_low','spearman_ci_high'])
        for r in res:w.writerow([r['system'],r['n'],r['pearson_r'],r['pearson_perm_p'],*r['pearson_bootstrap_95'],r['spearman_rho'],r['spearman_perm_p'],*r['spearman_bootstrap_95']])
    print(DATA/'robustness_statistics.csv')
if __name__=='__main__':main()
```
