## Supplementary material for "Projection criteria and information risks for zero-dimensional biological dynamics across molecular, epidemic, and ecological systems": Python scripts used to reproduce the simulations, numerical analyses, parameter screens, and figures reported in the manuscript.: run_all.py

```
#!/usr/bin/env python3
"""Reproduce all main-text numerical data and figures in isolated processes.

Run from any directory. Each script is launched in a fresh Python process to
avoid cross-script random-state, threading, or library-state interactions.
Runtime is machine dependent.
"""
from __future__ import annotations
from pathlib import Path
import os
import subprocess
import sys

HERE = Path(__file__).resolve().parent
ROOT = HERE.parent
env = os.environ.copy()
env.setdefault('OMP_NUM_THREADS', '1')
env.setdefault('OPENBLAS_NUM_THREADS', '1')
env.setdefault('MKL_NUM_THREADS', '1')

commands = [
    [sys.executable, str(HERE/'local_exchange_sensitivity.py')],
    [sys.executable, str(HERE/'spatial_sir_benchmark.py')],
    [sys.executable, str(HERE/'spatial_lv_benchmark.py')],
    [sys.executable, str(HERE/'two_compartment_model.py'), '--out', str(ROOT/'data'/'two_compartment_summary.csv')],
    [sys.executable, str(HERE/'generate_fig1_projection.py')],
    [sys.executable, str(HERE/'generate_fig2_survival.py')],
    [sys.executable, str(HERE/'generate_fig3_error.py')],
    [sys.executable, str(HERE/'generate_fig4_predictive_kl.py')],
    [sys.executable, str(HERE/'generate_fig5_parameter_map.py')],
    [sys.executable, str(HERE/'generate_fig6_ecoli_window.py')],
    [sys.executable, str(HERE/'generate_cross_scale_figures.py')],
    [sys.executable, str(HERE/'robustness_statistics.py')],
]

for command in commands:
    print('+', ' '.join(command), flush=True)
    subprocess.run(command, cwd=ROOT, env=env, check=True)

print('All benchmark data, robustness analyses, and figures generated.', flush=True)
```
