## Supplementary material for "Projection criteria and information risks for zero-dimensional biological dynamics across molecular, epidemic, and ecological systems": Python scripts used to reproduce the simulations, numerical analyses, parameter screens, and figures reported in the manuscript.: screen_0d_validity.py

```
#!/usr/bin/env python3
"""Screen spatial mixing and copy-number conditions for a 0D biochemical model.

Units:
  concentration: mol/L (M)
  volume: femtoliters (fL); 1 fL = 1 micrometre^3
  relative diffusion coefficient: micrometre^2/s
  bimolecular rate constant: M^-1 s^-1

The thresholds are user-selected diagnostics, not universal error guarantees.
"""
from __future__ import annotations
import argparse
import json
import math
from pathlib import Path

AVOGADRO = 6.02214076e23


def screen(concentration_M: float, volume_fL: float, Drel_um2_s: float,
           k2_M_inv_s: float, noise_tolerance: float = 0.1,
           da_tolerance: float = 0.1) -> dict[str, float | str | bool]:
    vals = [concentration_M, volume_fL, Drel_um2_s, k2_M_inv_s,
            noise_tolerance, da_tolerance]
    if any(x <= 0 for x in vals):
        raise ValueError("All inputs and tolerances must be positive.")
    L_um = volume_fL ** (1.0 / 3.0)
    pseudo_rate_s = k2_M_inv_s * concentration_M
    tau_mix_s = L_um**2 / (math.pi**2 * Drel_um2_s)
    tau_rxn_s = 1.0 / pseudo_rate_s
    da = tau_mix_s / tau_rxn_s
    copy_number = AVOGADRO * concentration_M * volume_fL * 1.0e-15
    rel_noise = 1.0 / math.sqrt(copy_number)
    N_required = noise_tolerance ** -2
    Vmin_fL = N_required / (AVOGADRO * concentration_M * 1.0e-15)
    Vmax_fL = (da_tolerance * math.pi**2 * Drel_um2_s / pseudo_rate_s) ** 1.5
    if da < 0.1:
        spatial_band = "rapid-mixing screen"
    elif da <= 1.0:
        spatial_band = "intermediate; validate spatially"
    else:
        spatial_band = "spatial-model warning"
    return {
        "concentration_M": concentration_M,
        "volume_fL": volume_fL,
        "length_um": L_um,
        "Drel_um2_s": Drel_um2_s,
        "k2_M_inv_s": k2_M_inv_s,
        "pseudo_first_order_rate_s_inv": pseudo_rate_s,
        "tau_mix_s": tau_mix_s,
        "tau_rxn_s": tau_rxn_s,
        "spatial_Damkohler": da,
        "spatial_screen": spatial_band,
        "mean_copy_number": copy_number,
        "poisson_relative_noise": rel_noise,
        "noise_tolerance": noise_tolerance,
        "copy_number_requirement": N_required,
        "da_tolerance": da_tolerance,
        "volume_lower_fL": Vmin_fL,
        "volume_upper_fL": Vmax_fL,
        "selected_volume_window_nonempty": Vmin_fL < Vmax_fL,
        "current_point_meets_selected_screens": rel_noise <= noise_tolerance and da <= da_tolerance,
    }


def main() -> None:
    ap = argparse.ArgumentParser(description=__doc__)
    ap.add_argument("--c", type=float, required=True, help="partner concentration in M")
    ap.add_argument("--V", type=float, required=True, help="volume in fL")
    ap.add_argument("--Drel", type=float, required=True, help="relative diffusivity in um^2/s")
    ap.add_argument("--k2", type=float, required=True, help="bimolecular rate in M^-1 s^-1")
    ap.add_argument("--noise-tol", type=float, default=0.1, help="selected relative-noise tolerance")
    ap.add_argument("--da-tol", type=float, default=0.1, help="selected Damkohler tolerance")
    ap.add_argument("--out", type=Path, help="optional JSON output path")
    args = ap.parse_args()
    result = screen(args.c, args.V, args.Drel, args.k2, args.noise_tol, args.da_tol)
    text = json.dumps(result, indent=2, ensure_ascii=True)
    print(text)
    if args.out:
        args.out.parent.mkdir(parents=True, exist_ok=True)
        args.out.write_text(text + "\n", encoding="utf-8")


if __name__ == "__main__":
    main()
```
