## Supplementary material for "Projection criteria and information risks for zero-dimensional biological dynamics across molecular, epidemic, and ecological systems": Python scripts used to reproduce the simulations, numerical analyses, parameter screens, and figures reported in the manuscript.: spatial_lv_benchmark.py

```
#!/usr/bin/env python3
"""Spatial predator-prey contact-process benchmark on a periodic lattice."""
from __future__ import annotations
from pathlib import Path
import csv, math
import numpy as np
ROOT=Path(__file__).resolve().parents[1]; DATA=ROOT/'data'; DATA.mkdir(exist_ok=True)

def neigh(a): return (np.roll(a,1,0),np.roll(a,-1,0),np.roll(a,1,1),np.roll(a,-1,1))

def pair_mi(s,n_states=3):
    pairs=[]
    for b in (np.roll(s,-1,0),np.roll(s,-1,1)): pairs.append(np.stack([s.ravel(),b.ravel()],1))
    p=np.concatenate(pairs); j=np.zeros((n_states,n_states))
    for x,y in p:j[int(x),int(y)]+=1
    j/=j.sum(); px=j.sum(1); py=j.sum(0); z=0.0
    for x in range(n_states):
      for y in range(n_states):
        if j[x,y]>0 and px[x]>0 and py[y]>0:z+=j[x,y]*math.log(j[x,y]/(px[x]*py[y]))
    return z

def pred_intensity(s,a):
    prey_neigh=sum((x==1).astype(int) for x in neigh(s))
    return float(a*np.sum((s==2)*prey_neigh)/4.0)

def wm_intensity(s,a):
    x=np.mean(s==1); y=np.mean(s==2); return a*x*y*s.size

def pkl(l,m):
    if l<=0:return max(m,0.0)
    if m<=0:return float('inf')
    return l*math.log(l/m)-l+m

def init(L,mode,rng):
    s=np.zeros((L,L),np.int8)
    if mode=='mixed':
        u=rng.random((L,L)); s[u<0.45]=1; s[(u>=0.45)&(u<0.60)]=2
    elif mode=='segregated':
        s[:,:L//2]=1; s[:,L//2:]=0
        mask=rng.random((L,L//2))<0.30; s[:,L//2:][mask]=2
        holes=rng.random((L,L))<0.15; s[holes]=0
    else: raise ValueError(mode)
    return s

def move(s,m,dt,rng):
    q=1.0-math.exp(-m*dt)
    k=int(rng.binomial(s.size,q))
    if k>1:
        idx=rng.choice(s.size,k,replace=False)
        vals=s.ravel()[idx].copy();rng.shuffle(vals);s.ravel()[idx]=vals

def sim(L=28,b=1.0,a=2.0,d=0.55,m=1.0,mode='mixed',T=25,dt=0.05,seed=0):
    rng=np.random.default_rng(seed); s=init(L,mode,rng); steps=int(T/dt); save=4
    tt=[]; tr=[]; risks=[]; kls=[]
    for step in range(steps+1):
      if step%save==0:
        tt.append(step*dt); tr.append([np.mean(s==0),np.mean(s==1),np.mean(s==2)])
        risks.append(pair_mi(s)); kls.append(pkl(pred_intensity(s,a),wm_intensity(s,a))/s.size)
      if step==steps:break
      move(s,m,dt,rng)
      prey_n=sum((x==1).astype(int) for x in neigh(s))
      pred_n=sum((x==2).astype(int) for x in neigh(s))
      birth=(s==0)&(rng.random(s.shape)<(1.0-np.exp(-b*prey_n*dt/4.0)))
      eaten=(s==1)&(rng.random(s.shape)<(1.0-np.exp(-a*pred_n*dt/4.0)))
      death=(s==2)&(rng.random(s.shape)<(1.0-math.exp(-d*dt)))
      s[death]=0; s[birth]=1; s[eaten]=2
    return np.array(tt),np.array(tr),np.array(risks),np.array(kls)

def ode(b,a,d,y0,t):
    y=np.array(y0,float);out=[y.copy()]
    def f(v):
      e,x,p=v; birth=b*x*e; pred=a*x*p; death=d*p
      return np.array([death-birth,birth-pred,pred-death])
    for t0,t1 in zip(t[:-1],t[1:]):
      h=t1-t0;k1=f(y);k2=f(y+h*k1/2);k3=f(y+h*k2/2);k4=f(y+h*k3)
      y=y+h*(k1+2*k2+2*k3+k4)/6;y=np.maximum(y,0);y/=max(y.sum(),1e-12);out.append(y.copy())
    return np.array(out)

def main():
    b=1.0;a=2.0;d=0.55;L=24;reps=8;ms=[0.0,0.2,1.0,5.0]
    rows=[];curves=[]
    for mode in ['mixed','segregated']:
      for m in ms:
        dat=[];ris=[];kls=[]
        for r in range(reps):
          t,tr,ri,kl=sim(L=L,b=b,a=a,d=d,m=m,mode=mode,seed=30000+r+int(100*m)+(0 if mode=='mixed' else 5000))
          dat.append(tr);ris.append(ri);kls.append(kl)
        mean=np.mean(dat,0);sd=np.std(dat,0);od=ode(b,a,d,mean[0],t)
        rmse=float(np.sqrt(np.mean((mean[:,1:]-od[:,1:])**2)))
        risk=float(np.mean(np.mean(ris,0)[:max(2,len(t)//3)]));ratekl=float(np.mean(np.mean(kls,0)[:max(2,len(t)//3)]))
        ext=float(np.mean([x[-1,2]<1/L**2 for x in dat]))
        rows.append([mode,m,risk,ratekl,rmse,ext,float(mean[:,1].max()),float(mean[:,2].max())])
        for ti,v,sv,ov in zip(t,mean,sd,od):curves.append([mode,m,ti,*v,*sv,*ov])
    with (DATA/'lv_benchmark_summary.csv').open('w',newline='') as f:
      w=csv.writer(f);w.writerow(['initial_geometry','mobility','pair_KL_nats','rate_KL_per_site','density_RMSE','predator_extinction_fraction','max_prey','max_predator']);w.writerows(rows)
    with (DATA/'lv_benchmark_curves.csv').open('w',newline='') as f:
      w=csv.writer(f);w.writerow(['initial_geometry','mobility','time','empty_mean','prey_mean','predator_mean','empty_sd','prey_sd','predator_sd','empty_ode','prey_ode','predator_ode']);w.writerows(curves)
    print(DATA/'lv_benchmark_summary.csv')
if __name__=='__main__':main()
```
