## Supplementary material for "Projection criteria and information risks for zero-dimensional biological dynamics across molecular, epidemic, and ecological systems": Python scripts used to reproduce the simulations, numerical analyses, parameter screens, and figures reported in the manuscript.: spatial_sir_benchmark.py

```
#!/usr/bin/env python3
"""Spatial SIR benchmark on a periodic square lattice.

The count projection retains only (S,I,R). Local infection depends on SI
nearest-neighbour contacts, so count-equivalent spatial configurations need
not have the same aggregate infection rate.
"""
from __future__ import annotations
from pathlib import Path
import csv, math
import numpy as np

ROOT = Path(__file__).resolve().parents[1]
DATA = ROOT / "data"
DATA.mkdir(exist_ok=True)


def neighbours(a: np.ndarray) -> tuple[np.ndarray, ...]:
    return (np.roll(a,1,0), np.roll(a,-1,0), np.roll(a,1,1), np.roll(a,-1,1))


def pair_mutual_information(state: np.ndarray, n_states: int=3) -> float:
    pairs=[]
    for b in (np.roll(state,-1,0), np.roll(state,-1,1)):
        pairs.append(np.stack([state.ravel(), b.ravel()], axis=1))
    p=np.concatenate(pairs, axis=0)
    joint=np.zeros((n_states,n_states),float)
    for i,j in p: joint[int(i),int(j)] += 1
    joint /= joint.sum()
    pi=joint.sum(axis=1); pj=joint.sum(axis=0)
    out=0.0
    for i in range(n_states):
        for j in range(n_states):
            if joint[i,j] > 0 and pi[i] > 0 and pj[j] > 0:
                out += joint[i,j]*math.log(joint[i,j]/(pi[i]*pj[j]))
    return out


def infection_intensity(state: np.ndarray, beta: float) -> float:
    infected_neigh=sum((x==1).astype(int) for x in neighbours(state))
    return float(beta*np.sum((state==0)*infected_neigh)/4.0)


def wellmixed_intensity(state: np.ndarray, beta: float) -> float:
    S=float(np.sum(state==0)); I=float(np.sum(state==1)); N=state.size
    return beta*S*I/N


def poisson_kl(lam: float, mu: float) -> float:
    if lam <= 0: return max(mu,0.0)
    if mu <= 0: return float("inf")
    return lam*math.log(lam/mu)-lam+mu


def init_state(L: int, frac_i: float, mode: str, rng: np.random.Generator) -> np.ndarray:
    n=L*L; ni=max(1,int(round(frac_i*n)))
    state=np.zeros((L,L),dtype=np.int8)
    if mode=="mixed":
        idx=rng.choice(n,ni,replace=False); state.ravel()[idx]=1
    elif mode=="clustered":
        cx=cy=L//2
        coords=[(i,j) for i in range(L) for j in range(L)]
        coords.sort(key=lambda z:(z[0]-cx)**2+(z[1]-cy)**2)
        for i,j in coords[:ni]: state[i,j]=1
    else: raise ValueError(mode)
    return state


def mobility_swaps(state: np.ndarray, mobility: float, dt: float, rng: np.random.Generator) -> None:
    q=1.0-math.exp(-mobility*dt)
    k=int(rng.binomial(state.size,q))
    if k>1:
        idx=rng.choice(state.size,k,replace=False)
        vals=state.ravel()[idx].copy(); rng.shuffle(vals); state.ravel()[idx]=vals

def simulate(L=24,beta=1.5,gamma=0.4,mobility=1.0,mode="mixed",T=20.0,dt=0.05,seed=0):
    rng=np.random.default_rng(seed); state=init_state(L,0.04,mode,rng)
    steps=int(round(T/dt)); save_every=2
    times=[]; traj=[]; risks=[]; ratekls=[]
    for step in range(steps+1):
        if step%save_every==0:
            times.append(step*dt)
            traj.append([np.mean(state==0),np.mean(state==1),np.mean(state==2)])
            risks.append(pair_mutual_information(state))
            la=infection_intensity(state,beta); lm=wellmixed_intensity(state,beta)
            ratekls.append(poisson_kl(la,lm)/max(state.size,1))
        if step==steps: break
        mobility_swaps(state,mobility,dt,rng)
        inf_neigh=sum((x==1).astype(int) for x in neighbours(state))
        p_inf=1.0-np.exp(-beta*inf_neigh*dt/4.0)
        p_rec=1.0-math.exp(-gamma*dt)
        infect=(state==0)&(rng.random(state.shape)<p_inf)
        recover=(state==1)&(rng.random(state.shape)<p_rec)
        state[recover]=2; state[infect]=1
    return np.array(times),np.array(traj),np.array(risks),np.array(ratekls)


def ode(beta,gamma,y0,times):
    y=np.array(y0,float); out=[y.copy()]
    def f(v):
        s,i,r=v; return np.array([-beta*s*i,beta*s*i-gamma*i,gamma*i])
    for a,b in zip(times[:-1],times[1:]):
        h=b-a
        k1=f(y); k2=f(y+h*k1/2); k3=f(y+h*k2/2); k4=f(y+h*k3)
        y=y+h*(k1+2*k2+2*k3+k4)/6
        y=np.maximum(y,0); y/=y.sum(); out.append(y.copy())
    return np.array(out)


def main():
    beta=1.5; gamma=0.4; L=20; reps=10
    mobilities=[0.0,0.2,1.0,5.0]
    rows=[]; curves=[]
    for mode in ["mixed","clustered"]:
        for m in mobilities:
            reps_data=[]; reps_r=[]; reps_k=[]
            for r in range(reps):
                t,tr,ri,kl=simulate(L=L,beta=beta,gamma=gamma,mobility=m,mode=mode,seed=10000+r+int(100*m)+(0 if mode=='mixed' else 5000))
                reps_data.append(tr); reps_r.append(ri); reps_k.append(kl)
            mean=np.mean(reps_data,axis=0); sd=np.std(reps_data,axis=0)
            od=ode(beta,gamma,mean[0],t)
            rmse=float(np.sqrt(np.mean((mean[:,1]-od[:,1])**2)))
            final_err=float(abs(mean[-1,2]-od[-1,2]))
            risk=float(np.mean(np.mean(reps_r,axis=0)[:max(2,len(t)//3)]))
            ratekl=float(np.mean(np.mean(reps_k,axis=0)[:max(2,len(t)//3)]))
            rows.append([mode,m,risk,ratekl,rmse,final_err,float(mean[:,1].max()),float(t[np.argmax(mean[:,1])])])
            for ti,v,sv,ov in zip(t,mean,sd,od): curves.append([mode,m,ti,*v,*sv,*ov])
    with (DATA/'sir_benchmark_summary.csv').open('w',newline='') as f:
        w=csv.writer(f); w.writerow(['initial_geometry','mobility','pair_KL_nats','rate_KL_per_site','infected_RMSE','final_size_error','peak_infected','peak_time']); w.writerows(rows)
    with (DATA/'sir_benchmark_curves.csv').open('w',newline='') as f:
        w=csv.writer(f); w.writerow(['initial_geometry','mobility','time','S_mean','I_mean','R_mean','S_sd','I_sd','R_sd','S_ode','I_ode','R_ode']); w.writerows(curves)
    print(DATA/'sir_benchmark_summary.csv')

if __name__=='__main__': main()
```
