## Supplementary material for "Projection criteria and information risks for zero-dimensional biological dynamics across molecular, epidemic, and ecological systems": Python scripts used to reproduce the simulations, numerical analyses, parameter screens, and figures reported in the manuscript.: two_compartment_model.py

```
#!/usr/bin/env python3
"""Exact benchmark for a hidden two-compartment bimolecular reaction.

One A and one B molecule occupy either the same compartment (C) or
separate compartments (S). Each molecule switches compartment at rate d,
so C <-> S occurs at total rate 2d. Reaction occurs from C at rate k.
The count projection cannot distinguish C and S.
"""
from __future__ import annotations

import argparse
from pathlib import Path
import numpy as np


def decay_rates(k: float, d: float) -> tuple[float, float]:
    if k <= 0 or d < 0:
        raise ValueError("Require k>0 and d>=0")
    disc = np.sqrt(k * k + 16.0 * d * d)
    alpha_minus = 0.5 * (k + 4.0 * d - disc)
    alpha_plus = 0.5 * (k + 4.0 * d + disc)
    return alpha_minus, alpha_plus


def survival(t: np.ndarray, k: float, d: float, p_colocated: float) -> np.ndarray:
    if not 0.0 <= p_colocated <= 1.0:
        raise ValueError("p_colocated must be in [0,1]")
    if d == 0.0:
        return (1.0 - p_colocated) + p_colocated * np.exp(-k * t)
    am, ap = decay_rates(k, d)
    gap = ap - am
    a = (ap - k * p_colocated) / gap
    b = (k * p_colocated - am) / gap
    return a * np.exp(-am * t) + b * np.exp(-ap * t)


def hazard(t: np.ndarray, k: float, d: float, p_colocated: float) -> np.ndarray:
    if d == 0.0:
        s = survival(t, k, d, p_colocated)
        return np.divide(k * p_colocated * np.exp(-k * t), s, out=np.zeros_like(t), where=s>0)
    am, ap = decay_rates(k, d)
    gap = ap - am
    a = (ap - k * p_colocated) / gap
    b = (k * p_colocated - am) / gap
    s = a * np.exp(-am * t) + b * np.exp(-ap * t)
    num = am * a * np.exp(-am * t) + ap * b * np.exp(-ap * t)
    return num / s


def main() -> None:
    parser = argparse.ArgumentParser()
    parser.add_argument("--out", default="../data/two_compartment_summary.csv")
    parser.add_argument("--k", type=float, default=1.0)
    parser.add_argument("--ratios", type=float, nargs="+", default=[0.1, 1.0, 10.0, 100.0])
    args = parser.parse_args()
    out = Path(args.out)
    out.parent.mkdir(parents=True, exist_ok=True)
    rows = ["d_over_k,alpha_minus_over_k,alpha_plus_over_k,max_error_equilibrium_init"]
    t = np.linspace(0.0, 10.0 / args.k, 4001)
    ref = np.exp(-0.5 * args.k * t)
    for ratio in args.ratios:
        d = ratio * args.k
        am, ap = decay_rates(args.k, d)
        s = survival(t, args.k, d, 0.5)
        err = float(np.max(np.abs(s - ref)))
        rows.append(f"{ratio:.12g},{am/args.k:.12g},{ap/args.k:.12g},{err:.12g}")
    out.write_text("\n".join(rows) + "\n", encoding="utf-8")
    print(out)


if __name__ == "__main__":
    main()
```
