## Supplementary material for "Projection criteria and information risks for zero-dimensional biological dynamics across molecular, epidemic, and ecological systems": Machine-readable CSV and JSON data underlying the numerical results and figures.: README_DATA.txt

Machine-readable data underlying the numerical results.

Main benchmark outputs:
- sir_benchmark_summary.csv, sir_benchmark_curves.csv
- lv_benchmark_summary.csv, lv_benchmark_curves.csv
- cross_scale_points.csv

Robustness outputs:
- sir_local_exchange_summary.csv
- lv_local_exchange_summary.csv
- local_exchange_correlations.csv
- robustness_statistics.csv and robustness_statistics.json

Molecular and screening outputs:
- two_compartment_summary.csv
- fig2_survival_data.csv through fig6_ecoli_window_data.csv
- ecoli_case_summary.csv and JSON screen files

All files are generated by scripts in S1 Code. Random seeds and parameter values are embedded in the scripts and described in the manuscript and Supplementary Material.
